# Bridging the light-electron resolution gap with correlative cryo-SRRF and dual-axis cryo-STEM tomography

**DOI:** 10.1101/2022.11.19.517200

**Authors:** Peter Kirchweger, Debakshi Mullick, Prabhu Prasad Swain, Sharon G. Wolf, Michael Elbaum

**Affiliations:** Department of Chemical and Structural Biology, Weizmann Institute of Science, 7610001 Rehovot, Israel; Department of Chemical and Biological Physics, Weizmann Institute of Science, 7610001 Rehovot, Israel; Diamond Light Source: Didcot, Oxfordshire, GB; School of Physical Sciences, UM-DAE Centre for Excellence in Basic Sciences, Mumbai 400098, India; Institute of Bioengineering, Swiss Federal Institute of Technology Lausanne (EPFL), CH-1015, Lausanne, Switzerland; Department of Chemical Research Support, Weizmann Institute of Science, 7610001 Rehovot, Israel

## Abstract

Cryo-electron tomography (cryo-ET) is the prime method for cell biological studies in three dimensions (3D) at high resolution. We have introduced cryo-scanning transmission electron tomography (CSTET), which can access larger 3D volumes, on the scale of 1 micron, making it ideal to study organelles and their interactions *in situ*. Here we introduce two relevant advances: a) we demonstrate the utility of multi-color super-resolution radial fluctuation light microscopy under cryogenic conditions (cryo-SRRF), and b) we extend the use of deconvolution processing for dual-axis CSTET data. We show that cryo-SRRF nanoscopy is able to reach resolutions in the range of 100 nm, using commonly available fluorophores and a conventional widefield microscope for cryo-correlative light-electron microscopy (cryo-CLEM). Such resolution aids in precisely identifying regions of interest before tomographic acquisition and enhances precision in localizing features of interest within the 3D reconstruction. Dual-axis CSTET tilt series data and application of entropy regularized deconvolution during post-processing results in close-to isotropic resolution in the reconstruction without averaging. We show individual protein densities in a mitochondrion-ER contact in a cell region 850 nm thick. The integration of cryo-SRRF with deconvolved dual-axis CSTET provides a versatile workflow for studying unique objects in a cell.

## 1 Introduction

Cryo-transmission electron tomography (cryo-ET, [1]), cryo-scanning transmission electron tomography (CSTET, [2]), cryo-soft X-ray tomography (cSXT, [3]), and cryo-focused ion beam-scanning electron microscopy (cryo-FIB-SEM, [4–7]) are complementary methods to visualize unstained vitrified cells in three-dimensions (3D) at different scales [8, 9]. Cryo-ET uses a parallel beam to record magnified projections of the region of interest on a camera and exploits phase contrast to achieve the highest resolutions [10]. CSTET, on the other hand, scans a focused electron beam across the region of interest (Fig S1A) while scattered electrons are recorded on area detectors. CSTET is able to image thicker specimens than conventional cryo-ET, albeit at a lower resolution [2, 11–13]. cSXT takes advantage of the “water window” between the carbon and oxygen K-absorption edges in the electromagnetic spectrum; organic matter then appears dark on the relatively transparent aqueous background. It is normally practiced at synchrotron facilities, although suitable lab sources are emerging (e.g. [14]. All three have in common the Radon scheme of reconstruction from tilted projections [15]. Cryo-FIB-SEM works by an entirely different principle based on iterative ablation of a thin layer of the specimen and then imaging of the freshly exposed surface.

In tilt-based tomographies, a series of images are recorded as the specimen is rotated around an axis. The images are then aligned to compensate for mechanical or optical distortions. The final 3D volume is reconstructed by one of the available algorithms, which include weighted back-projection, Simultaneous Iterative Reconstruction Technique (SIRT), algebraic reconstruction, and direct Fourier inversion [16]. These produce similar but not identical results, as the complete 3D inversion problem is typically under-determined by the limited amount of 2D data available. Geometrical restrictions limit the tilt range to about 120°, which leads to a large “missing wedge” of information; discrete tilts also impose small missing wedges. Collecting a second tilt series at the same position by rotating the grid 90° reduces the missing wedge to a missing pyramid [17, 18] (Fig S1*B*). This method is known as dual-axis tomography and in principle, can be carried further to multiple axes [19]. Due to the increased radiation exposure, dual-axis tomography is mainly used for plastic sections [17, 18, 20, 21]). However, cryo-ET using a Volta Phase Plate was recently extended to dual-axis tomography for the study of Ebola virus-like particles [22].

While TEM-based cryo-ET has been instrumental in elucidating the morphology and ultra-structure of biological specimens, it is limited to very thin areas of the specimen. The specimen thickness limit arises mainly from inelastic scattering of the electrons. Chromatic aberration in the objective then produces a low-resolution haze in the image, as different energy losses come to focus in different planes [23]. An imaging spectrometer is normally used to select the electrons with close to zero energy loss, effectively removing the haze. However, the sample continues to suffer damage while the image signal is reduced. The physical limit for energy-filtered TEM tomography, therefore, relates to the inelastic mean free path of water, roughly 300 nm for 300 keV electrons [24]. To circumvent this issue, a focused ion beam (FIB) is often used to mill thin lamellae in thicker specimens, such as cells [25–27]. CSTET, on the other hand, is insensitive to chromatic aberration because there is no image-forming lens after the specimen. As such, no energy filter is required, and the dose efficiency is also higher for thicker samples. Loss of contrast reflects the onset of multiple elastic scattering; in practice, this permits a deep view into the cell on the order of one micron or more. This is similar to the thickness limit for vitrification by conventional plunging to liquid ethane. For example, intact mitochondria can be visualized in the context of surrounding cellular elements, such as the endoplasmic reticulum (ER) or microtubules (MT) [28], and in certain cells, even the edge of the nucleus is accessible [29].

Increased sample thickness accentuates the problem of sparse information for reconstruction. Artifacts of the reconstruction process are recognized as “ghost” contrast projecting into neighboring Z slices. In single-axis tomography, these are especially noticeable as streaks in XZ cuts of the volume emanating from high contrast features (e.g., gold nanoparticles), but they also create a “salt and pepper” noise of random appearance in distant XY planes. Given that the origin is structural, these artifacts can be suppressed algorithmically using prior knowledge of the 3D image of an idealized point object. This is implemented by deconvolution using a synthesized 3D point spread function (PSF). Entropy regularized deconvolution (ERDC) [30], developed originally for fluorescence microscopy, has been adapted specifically to this purpose [29, 31].

Another major challenge in cryo-tomography is to locate and identify the relevant areas of interest, both for acquisition and for analysis of the reconstruction. Correlative light-electron microscopy (cryo-CLEM, [32, 33]) is the most versatile tool available today, and specialized microscope stages are available for fluorescence imaging under cryogenic conditions. In practice, the final optical resolution constitutes a significant fraction of the entire field of view in a tomogram. Thus, there still remains a significant resolution gap between optical microscopy images and EM reconstruction.

Several attempts have been made to bridge this gap by implementing of high-resolution light imaging methods [7, 34–39]. Among these, many depend on specific microscopes, special fluorophores, or high intensity illumination; especially the last is challenging to use under cryogenic conditions.

In this work, we demonstrate a complete workflow involving cryo-CLEM with conventional fluorescence microscopy deconvolution (FMD) and high-resolution imaging based on super-resolution radial fluctuations [40, 41] under cryogenic conditions (cryo-SRRF). We imaged fibroblast cells grown directly on EM grids and stained with Hoechst, TMRE, SPY650-Tubulin, and BODIPY-ceramide for nucleus, mitochondria, microtubules, and the Golgi, respectively. We then collected dual-axis CSTET tilt series of intact cells, without sectioning, at specifically targeted regions of interest in areas 850 nm thick. Reconstructions were processed by ERDC [29] to minimize the missing wedge and pyramid effects. Combining ERDC with dual-axis tomography results in visually isotropic resolution also in the Z direction. Finally, the cryo-SRRF and fluorescence deconvolution were overlaid onto the CSTET volumes. We demonstrate and analyze the relevant improvements at each stage.

## 2 Materials and Methods

### 2.1 Cell culture and CSTET specimen preparation

Cells were grown as described in [42]. Briefly, WI38-lung fibroblasts were purchased from Coriell and maintained in Minimal Essential Medium (MEM), including 15% fetal bovine serum, L-glutamine, and penicillin/streptomycin according to the supplier’s instructions. The cells were seeded on EM grids (Quantifoil SiO2 R2/2, 200 mesh, gold) in a 3D printed specimen holder [43] and grown overnight.

The cells were labeled for Golgi, mitochondria, microtubules, and nucleus on the day of sample vitrification. Golgi was labeled with BODIPY FL C5-ceramide (B22650, ThermoFisher Scientific) according to the manufacture’s instructions. In short, the growth media was replaced with Hanks Buffered Salt Solution + 10 mM HEPES, pH 7.4 (HBSS/HEPES) containing 2.5 *μM* BODIPY. After 30 min at RT, the HBSS/HEPES was replaced with growth medium and incubated for an additional 30 min at 37°C, 5% CO2. Subsequently, mitochondria were labeled with 0.1 *μM* TMRE (T669, ThermoFisher Scientific) for 30 min in growth media. Microtubules were stained with SPY650-Tubulin (SC503, Spirochrome Probes). In short, a 1000x stock was prepared for storage, and for staining a 3x was added to the growth medium for 60 min. The nucleus was stained with 1 *μM* Hoechst (62249, ThermoFisher Scientific) for 5 min. Immediately before plunging, the medium containing the stains was replaced with fresh growth medium.

The grids were cryo-fixed in a Leica EM-GP plunger (Leica Microsystems, Vienna, Austria). The blotting chamber was set to 95% humidity at 37 °C, 2 μL of growth medium (front side of the grid), and 2 μL of 15 nm fiducial markers were added to the back side of the grid. The grids were blotted on the back side (away from the cells) for 5 sec, and plunged into liquid ethane after a post-blotting delay of 4 sec. Grids were stored in liquid nitrogen until use.

### 2.2 Cryo-FM grid map acquisition

The cryo-FM map was recorded in an EM Cryo CLEM (Leica Microsystems) as reported [44], except that the microscope is equipped with a Hamamatsu ORCA-Flash 4.0 instead of the standard camera. A focus map was recorded in the TexasRed (TXR) channel to focus on the mitochondria in the cells rather than the carbon. Subsequently, a full grid map was recorded using the BF and the TXR channel. A Z-stack of 15 *μm* in thickness with 20 slices was recorded, and single and combined mosaics were exported. Based on this map specific areas of interest were selected as potential candidates suitable for FMD, cryo-SRRF and CSTET data collection.

### 2.3 Cryo-FMD

For fluorescence microscopy deconvolution, an image stack of *∼*20*μm* was acquired at a Z-step of 300 nm in all four channels. Image analysis plugins available in FIJI [45] were used for all subsequent analysis. The image stack was first stabilized using the Image Stabilizer plugin [46]. A theoretical point spread function (PSF) was computed for each fluorescent wavelength with the PSF Generator plugin [47] using the classical Born and Wolf 3D optical model. The stabilized image stacks along with their matching PSFs were processed with the Deconvolution Lab2 plugin using a Richardson-Lucy algorithm with 200 iterations [48]. Maximum intensity projections from central slices were used for correlative microscopy.

### 2.4 Cryo-SRRF data acquisition and analysis

SRRF is a post-acquisition processing approach to obtain super-resolution fluorescence images based on a localization approach using radial symmetry [40, 41]. The novelty of this approach is the absence of a requirement for specific blinking of isolated fluorophores, exploiting instead intensity fluctuations that are largely unavoidable. This permits the use of any fluorescent dye and avoids the requirement for dye-specific buffers. Moreover, the use of low-intensity illumination circumvents concerns about sample heating and devitrification, making the method conducive for use at cryogenic temperatures. Using the Hamamatsu ORCA-Flash4.0 camera controlled by the Multi-Dimensional Acquisition module on μ-Manager [49, 50], we collected 500 frames of 1024×1024 pixels with a high frame rate (typically *∼*100 frames per second) at one Z-height. Only one plane with the sharpest focus was used, as the acquisition through the volume was challenging owing to drift issues. For the image analysis, we split the 500 frames into 5 stacks of 100 each and analyzed the 5 frames separately using the NanoJ plugin in FIJI [40, 41, 51]. Precise settings of the individual data sets appear in the supplementary information.

### 2.5 Cryo-SRRF and cryo-FMD edge response estimation

Single Z slices in each color channel were chosen at four representative positions, and a 2 *μm* long line was drawn in FIJI across these positions, and an intensity profile was measured along these lines. For calculating the edge response, [52] the maximum value of one-half of the intensity profile was chosen. The difference between the corresponding distances of 10% and 90% of the maximum value was calculated, which resulted in the edge response. The mean and standard deviation of these four points was calculated and is recorded in Table S1.

### 2.6 CSTET data collection

CSTET data of native vitrified cells were collected on a Titan Krios G3 (Thermo Fisher Scientific) equipped with an X-FEG electron source operated at 300 kV, a dual-axis tomography compustage and a Fischione high-angle annular dark-field detector (HAADF). Illumination conditions included a 70 *μm* condenser aperture, STEM probe semi-convergence angle of 1.2 mrad, and Spot size 9, which resulted in a current of 20 pA measured on the factory-calibrated Fluscreen. Bright Field (BF)-STEM data were collected on the HAADF detector by using a 20 *μm* objective aperture to limit the collection semi-angle to 3 mrad on the axis, and using the diffraction alignment to move the signal off-axis and onto the sensitive area of the detector.

SerialEM [53, 54] was used for data collection. A low magnification grid map (LM-STEM mode, 260x mag) was recorded to locate the cells of interest by correlation with the cryo-FM map. Medium magnification maps were acquired (microprobe mode, 3600k mag) for these cells, and then positions for tilt series data collection were chosen and anchor maps recorded. Tilt series were collected using the dose symmetric tilt scheme [55] up to +/-60° in 2° steps. Rough and fine eucentricity alignments were performed before setting up the position for data collection. Images of 2048 × 2048 pixels were recorded with a dwell time of 3 μsec/pixel at a magnification of 29000x for a pixel size of 2.1 nm/pixel. Dynamic focusing was activated to maintain focus across the tilted fields. Autofocusing was skipped due to the high depth of field in the CSTET data collection. With these parameters, a tilt series is collected in around 30 min. After recording the first axis on all desired positions, the grid was rotated in the microscope by 90°, each position was revisited manually, and the position was corrected with the “Realign to Item” option in SerialEM. The second axis was recorded using the same settings.

### 2.7 Tomogram reconstruction

Tilt series alignment and tomogram reconstruction were done using the batch reconstruction protocol of etomo in IMOD software (version 4.12.10; [56, 57]). Approximately 70 fiducials were used to align the tilt series automatically, followed by manual refinement. The tomograms were binned by 2 for visualization. Two reconstruction algorithms were used: a) standard weighted back-projection (WBP) for further deconvolution processing and b) SIRT-like filtered [58] for direct visualization and tomogram combination. More details about tomogram reconstruction and combination appear in the supplementary information. Final tomograms were visualized in UCSF ChimeraX [59, 60].

### 2.8 ERDC

ERDC of the dual-axis WBP CSTET reconstruction was adapted from [29] for dual axis and higher throughput. A Python script was written to perform most of the necessary steps (Fig S4, and Supplementary Methods). In short, a pre-calculated single-probe PSF, the dual-axis WBP tomogram, the refined tilt angles from both axes (in the form of the .tlt file (from IMOD) and the .patchcorr log file (from IMOD) were provided. The script first scales the tomogram and inverts the density from a black (high intensity)-on-white (low intensity) to a white-on-black contrast. It then generates the dual-axis multi-probe PSF by rotating the single-probe PSF according to the refined tilt angles (with rotatevol (IMOD)), summing them (with “clip add” (from IMOD)), and projecting the multi-probe PSF into Fourier space (using FTransform3D (Priism, [29, 30, 61])). Finally, the ERDC is performed on the tomogram using core2 decon (in Priism).

For diagnostic purposes, the script also provides summed power spectra of the input and output tomogram in the XY, XZ and YZ orientations using “clip spectrum”, followed by “clip avg” functions (IMOD).

Complete details about the usage of the Python script appear in the supplemental information.

### 2.9 Cryo-CLEM correlation

The FM images (source) were overlaid on the EM images (target) using the ec-CLEM plugin [62] in Icy [63]. The correlation is done by first overlapping the multi-channel fluorescence and BF images from the Leica cryo-CLEM microscope with the medium magnification EM overview (Anchor Map from SerialEM). Subsequently, the correlated image becomes a source for the target tomogram, which yields the final CLEM image. Mitochondria in the tomograms and their fluorescent signals were used as markers during the overlay, along with recognizable features from the BF channel.

### 2.10 IsoNet

We ran IsoNet (version 0.1, [64]) on the WBP and ERDC tomograms according to the tutorial instructions, with the exception of skipping CTF correction. The cube size was 96 voxels. 30 iterations were run with increasing noise correction from 0.05 to 0.2 in 4 steps. We visualized the resulting tomograms in UCSF ChimeraX [59, 60].

## 3 Results

### 3.1 Cryo-FM

Finding the object of interest in the feature-rich environment of an eukaryotic cell is a major challenge in cryo-ET. To address this, we correlated cryo-SRRF images with dual-axis CSTET. We collected a cryo-FM grid map of the plunge-frozen specimen with the bright field and TXR channel. Based on that map we chose cells of interest, for which we collected FMD, cryo-SRRF and dual-axis CSTET data. Subsequently, we reconstructed the CSTET tomograms of the same areas and processed them by ERDC (workflow diagram, Fig 1*A*).

**Fig 1:**
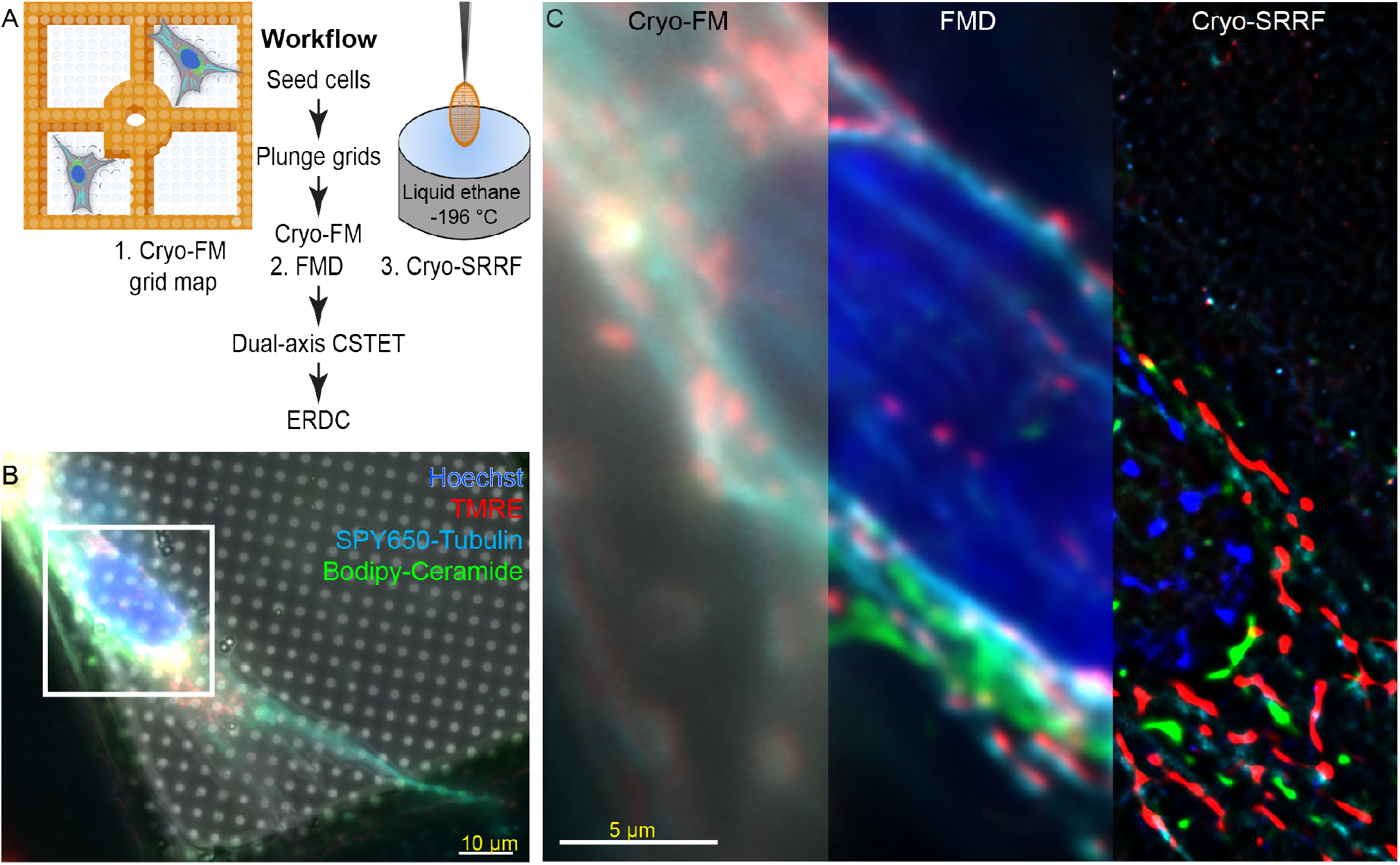
Workflow and Cryo-FM imaging: (A) Scheme of the workflow applied in this study. (B) Stabilized maxZ representation of the cryo-FM image without post-processing. Scale bar 10 *μm*. (C) Comparing stabilized maxZ to FMD and cryo-SRRF (the full images appear in Fig S2*A*). Blue, red, cyan, and green colors indicate heterochromatin, mitochondria, microtubules, and Golgi staining, respectively. Scale bar 5 *μm*.

The unprocessed, stabilized maximum-intensity Z-projection (maxZ) cryo-FM image reveals only very crude details about the stained organelles and cytoskeleton (Fig 1*B*). FMD reduces the haze in the maxZ image, but the lateral precision of the fluorescent signal remains low (Fig 1*C* (middle panel)). With cryo-SRRF, a much more precise localization is achieved (Fig 1*C* (right panel)). For the cryo-SRRF data sets, we obtained a consistent resolution on the order of 100-200 nm as estimated both by edge response functions and Fourier Ring Correlation (FRC). See Supplementary Table S1 and Fig S3. Specific values varied, with dependence on the fluorophore, the illumination intensity, and the camera exposure time. The improvement is highlighted by line scans of the cryo-SRRF signal in two areas in this cell (Fig S2*D,E*). The line scan peaks of the cryo-SRRF (Fig S2*D*, solid lines) are much sharper than those of the deconvolved cryo-FM data (Fig S2*D*, dotted lines).

The signal patterns and intensities from FMD and SRRF appear similar for mitochondria (Fig S2*D* (red plot)), but can be different for microtubules (Fig S2*D* (cyan plot)) and Golgi stain (Fig S2*D* (green plot)). While the FMD data originate from a Z stack extending through the entire cell, the cryo-SRRF signal was recorded only from a single plane and so represents an effective optical section.

### 3.2 Deconvolved dual-axis CSTET

#### 3.2.1 ERDC

Dual-axis tomograms were recorded in cell regions ranging from 700 nm (Fig S5) to 850 nm thickness (Fig 2). The field-of-view (FOV) for the chosen sampling (2 nm/pixel) is approximately 4.2 *μm*, allowing for observation of many intracellular interactions between mitochondria, ER, and microtubules.

Dual-axis WBP (Fig 2*A,D*) and SIRTlike30 filtered CSTET reconstructions (Fig 2*B,E*) reveal many ultrastructural details. Microtubules can be traced throughout the tomogram (cyan arrowheads). Two mitochondria, with diameters of about 400-450 nm, can be observed in proximity to ER. The organelles are 30-70 nm apart along a length of roughly 200 nm. A protein tether (white arrowhead) is seen at a contact site between the mitochondria and ER (red and green arrowheads, respectively) (Fig 2*D,E*), suggesting that this is indeed a bona fide mitochondria-ER contact [65]. However, at this level of representation, it is difficult to discern any further details.

Recently we described the entropy regularized deconvolution (ERDC) of CSTET data, which significantly enhances the quality of the tomograms by reducing the “salt and pepper” noise that results from the spurious projection of contrast into neighboring planes [29]. Here, we developed a more streamlined protocol for this method and extended it for application to dual-axis tomograms. In addition to in-plane noise reduction, ERDC improves the representation of objects in the axial direction, partly compensating for artifacts due to the missing wedge [29, 31]. The deconvolved tomogram reveals protein densities in the cristae and ER membranes (see red and green arrowheads, respectively (Fig 2*F*). These densities are present and visible in the SIRTlike30 filtered tomogram (Fig 2*E*), but the contrast is clearly enhanced by deconvolution. Additionally, several protein densities (white arrowheads) span the ER-mitochondria contact site at different distances. Ribosomes are also visible at the edge of the contact site (yellow arrowheads).

**Fig 2:**
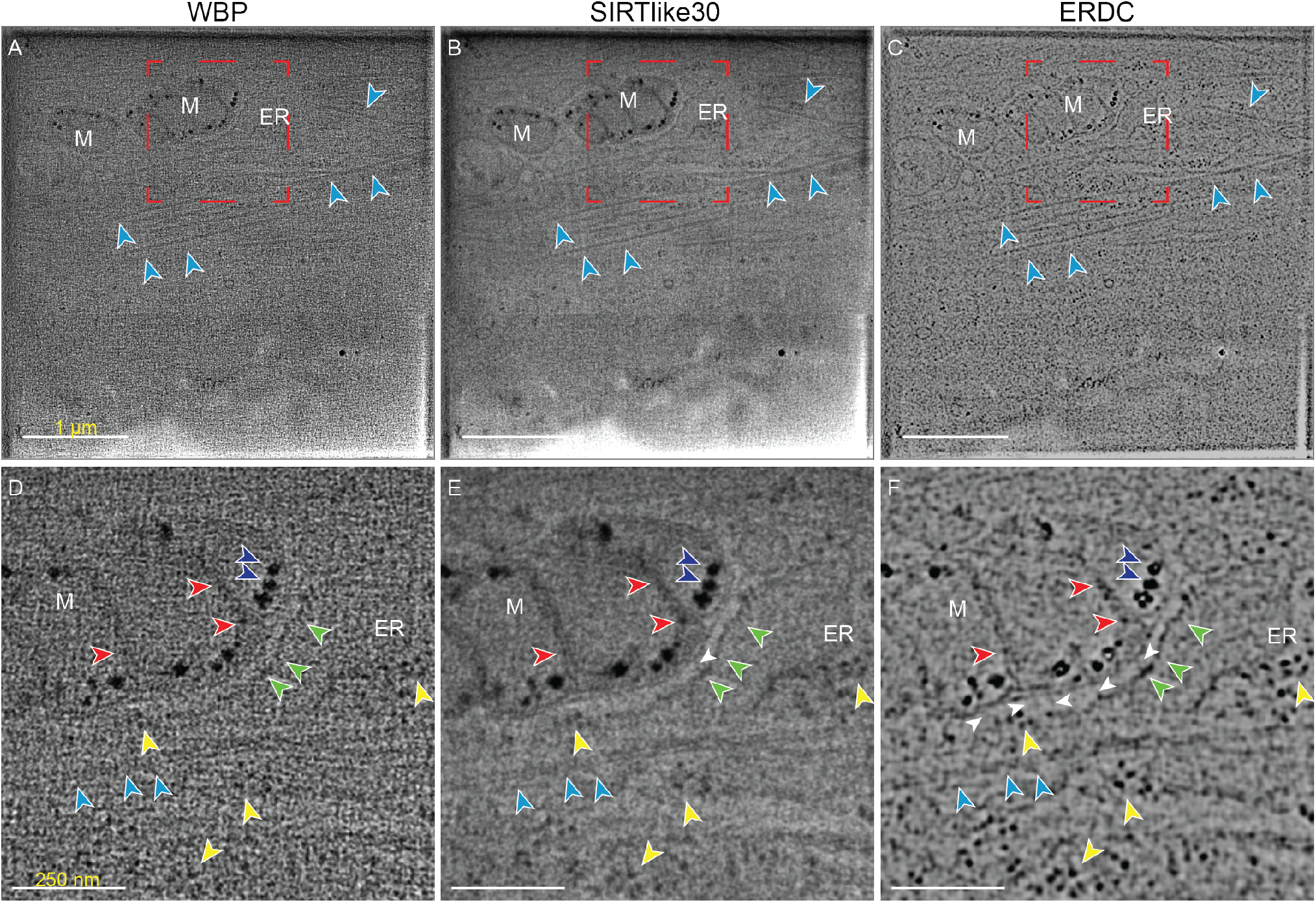
Dual-axis CSTET tomogram: Comparison of (A,D) WBP (input for deconvolution) and (B,E) SIRTlike30 filtered tomogram, reconstructed in IMOD, and (C,F) deconvolved dual-axis tomogram. (A,B,C) An overview of the whole tomogram. Scale bar 1 *μm*. (D,E,F) Close-ups showing contact between mitochondria (M) and ER (green arrowheads). Arrowheads point to cristae (red), Calcium phosphate deposits (blue), ER (green), ribosomes (yellow), microtubule (cyan), and protein densities in the mitochondria-ER contact area (white). Red box indicates the area depicted in the lower row.

While the noise suppression by ERDC enhances the quality of the tomogram in the XY plane, dramatic improvements are seen in the XZ and YZ planes. Two areas are chosen for comparing WBP, SIRTlike30 and ERDC (Fig 3). In the standard WBP many details are hidden due to the high level of noise (Fig 3*A,D*). Reconstruction with the SIRTlike30 filter suppresses some of the noise and reveals more details, such as a microtubule running horizontally to the XY plane (cyan arrowheads), the membranes of a vesicle (green arrowheads) (Fig 3*B*), and cristae and outer mitochondrial membrane (OMM) (red and orange arrowheads, respectively, Fig 3*E*). ERDC (Fig 3*C,F*) further enhances the contrast in the XZ and YZ planes. It reveals individual protein densities along the microtubules (cyan arrowheads), a pair of ribosomes (yellow arrowhead), and slices through mitochondria (orange and red arrowheads for the OMM and cristae, respectively)(Fig 3*C*). Most strikingly, protein densities are revealed on top of the mitochondria (Fig 3*F*, white arrowheads), where the missing wedge effect should be most severe. This demonstrates that ERDC indeed improves the quality of dual-axis tomograms, especially in the XZ and YZ planes.

**Fig 3:**
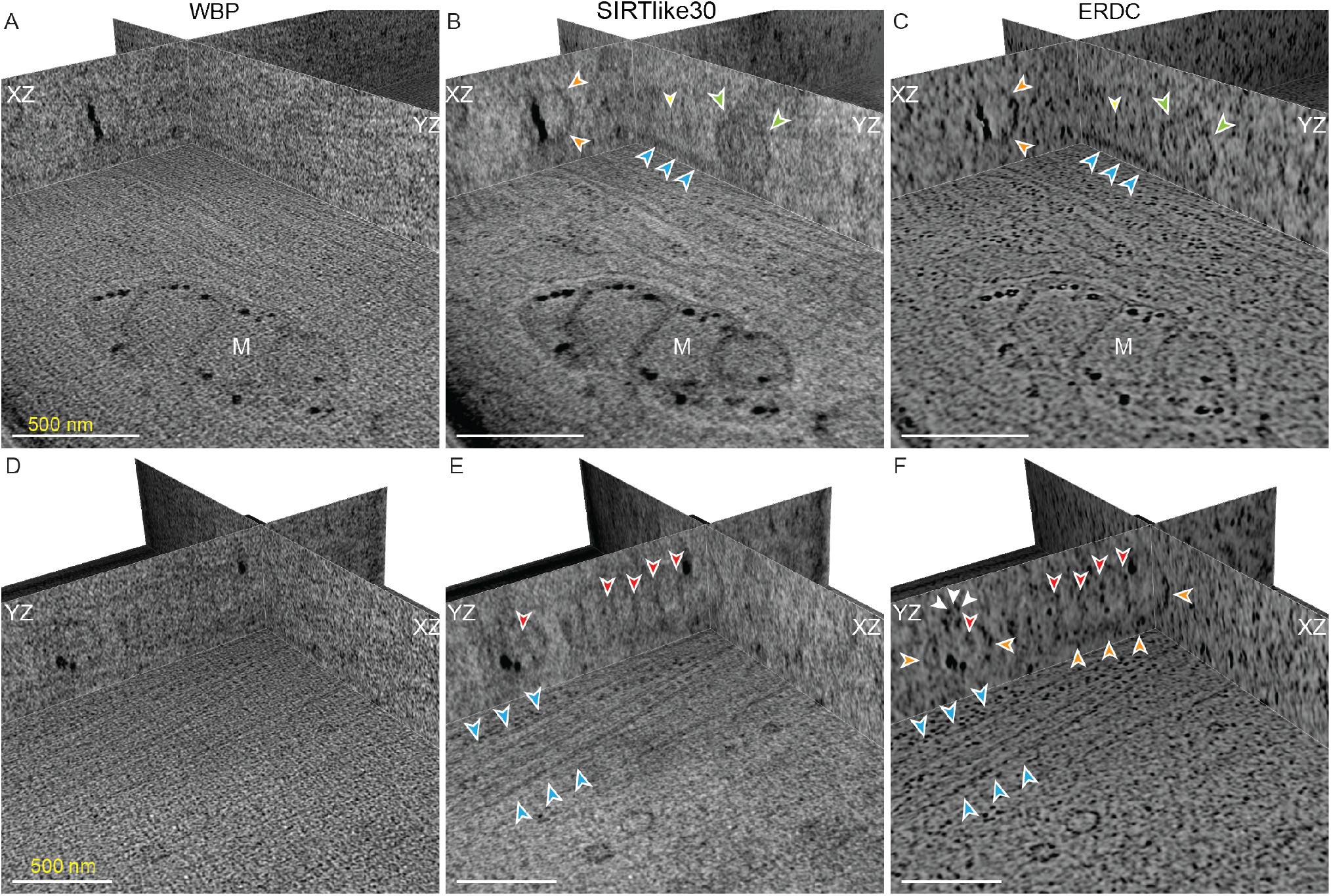
Orthoplane views of dual-axis CSTET tomograms: Comparison of (A,D) WBP (input for deconvolution) and (B,E) SIRTlike30 filtered tomogram, reconstructed in IMOD, and (C,F) deconvolved dual-axis tomograms. (A,B,C) A region showing microtubules (cyan arrowheads) running along the YZ plane, the membrane of a vesicle (green arrowheads), and ribosomes (yellow arrowheads). (D,E,F) A region showing the outline of cristae of mitochondria (red arrowheads) and the OMM (orange arrowheads). Scale bars are 500 nm. Total region thickness is 850 nm.

Volume rendering of 120 thick sections Fig 4 and Fig S6 with inverted density show many details. Ribosomes (yellow arrowheads) are seen as bright points, which often appear in spiral clusters as poly-ribosomes. Even in this thick section protein densities in the mitochondria-ER contact site (Fig S6, white arrowheads) are visible. Microtubules spanning the whole FOV can be traced, with individual protein densities becoming visible (cyan arrowheads). Due to the higher atomic number and stronger electron scattering, calcium phosphate granules inside the mitochondria are the most prominent feature (blue arrowhead).

**Fig 4:**
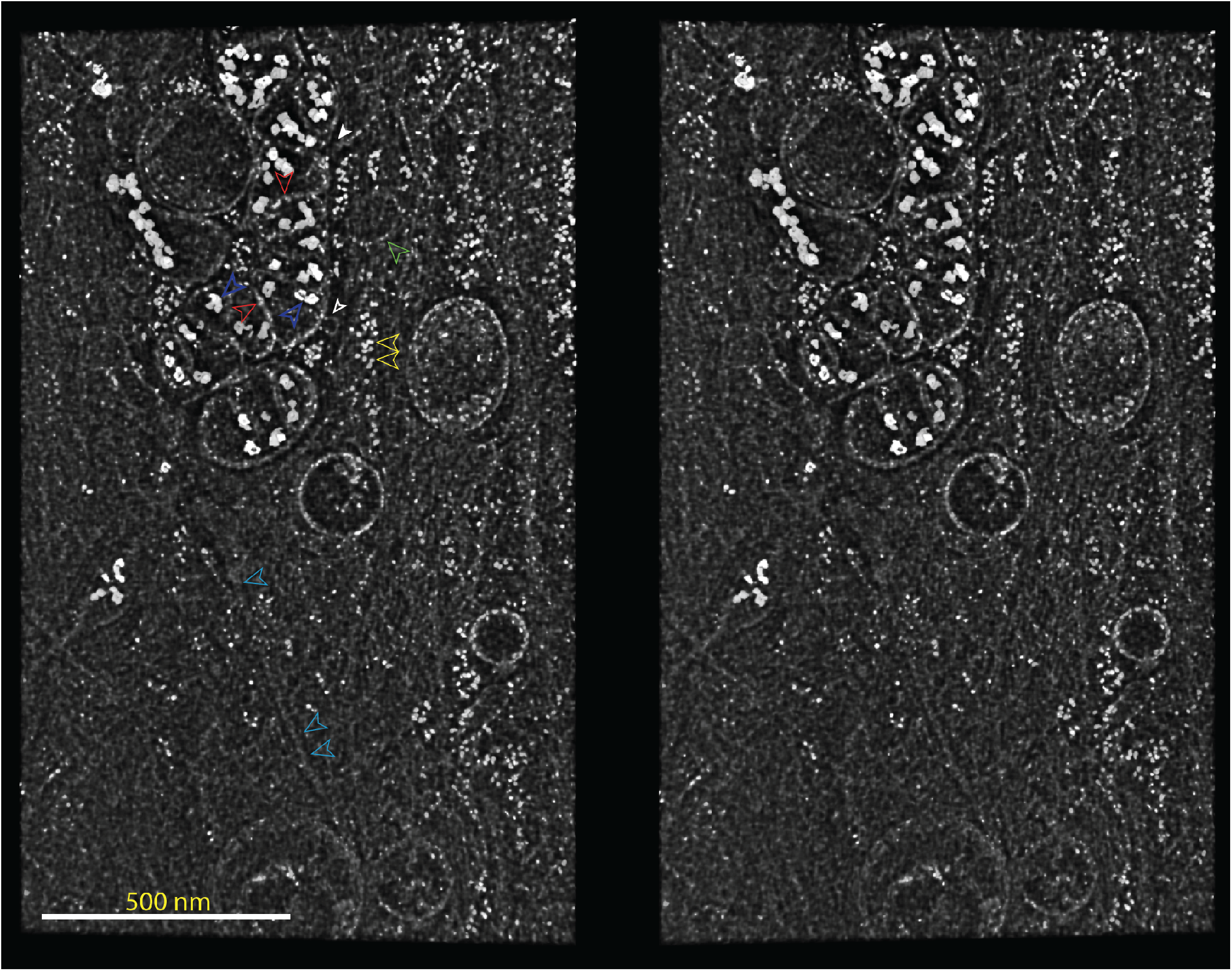
Stereo images of a 120 nm thick section of a different region in the deconvolved tomogram of Fig 2, with inverted contrast. Cristae (red arrowheads), calcium phosphate deposits (blue arrowheads), ER membrane (green arrowheads), ribosomes (yellow arrowheads), microtubules (cyan arrowheads), and unknown protein densities in contact with the mitochondria can be readily observed even in this thick section (white arrowheads). Slices 100-130 are depicted. Total region thickness is 850 nm.

#### 3.2.2 Comparing Single to Dual-axis tomography

Next, we compare the deconvolved dual-axis reconstruction shown in (Fig 3*C,F* with similar processing of one of the axes that produced it (Fig 5). As expected, the dual-axis tomograms reveal details that are missing in single-axis reconstructions, especially in the XZ and YZ planes. For example, the microtubule running in the YZ plane (cyan arrowheads, Fig 5*A*) is not seen in the single-axis version (Fig 5*B*), probably due to its orientation perpendicular to the tilt axis. Cristae (red arrowheads) are seen in both, (Fig 5*C,D*) but the OMM above and below the cristae are only seen in the dual-axis tomogram (orange arrowheads, Fig 5*C*, Fig S6).

**Fig 5:**
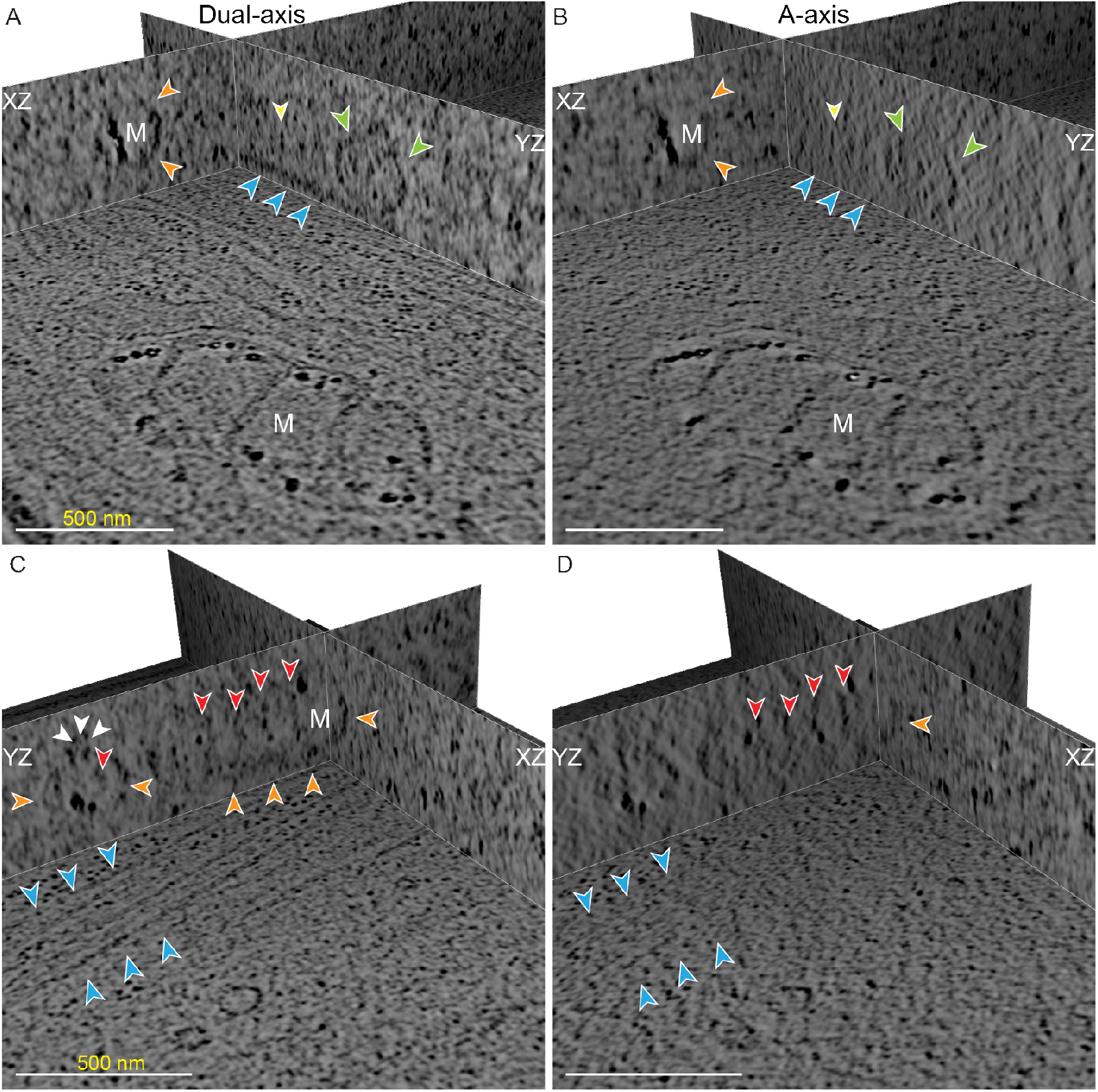
Comparing Dual- vs single-axis CSTET: Comparison of (A,C) deconvolved dual- and (B,D) single-axis tomogram. Orthoplanes of two areas in Fig 3 (indicated by the cristae (red arrowheads) and calcium phosphate deposits (blue arrowheads)) and ER (green arrowheads). Yellow arrowheads indicate ribosomes, cyan arrowheads point to microtubules, and white arrow-heads point to protein densities in the mitochondria-ER contact area. (A,C are the same images as Fig 3*C,F*)

### 3.3 Comparing ERDC to Deep Learning Approach

Next, we compare ERDC to IsoNet [64], which uses a neural network to generate a restoration filter for missing wedge data using sub-volumes in the reconstruction. A small region in the mitochondria-ER contact site is chosen for focused analysis (Fig 6*A*, white box) with XY, XZ, and YZ views displayed (second, third, and forth row, respectively). WBP does not reveal any details other than the prominent calcium phosphate deposits, especially in XZ and YZ (Fig 6*A*, blue arrowheads). In the SIRTlike30 reconstruction the XY planes show some details, such as the membrane of the ER, or a hint towards a protein tether (white arrowheads), but in the XZ and YZ views hardly anything can be discerned (Fig 6*B*). With ERDC, however, the density of the mitochondria-ER tether (white arrowheads) can be observed clearly in all views (white arrowheads) (Fig 6*C*). (A single white pixel is used to mark the identical location in all panels.) The OMM, the inner mitochondrial membrane, and the ER membrane can all be distinguished in the XZ and YZ planes.

**Fig 6:**
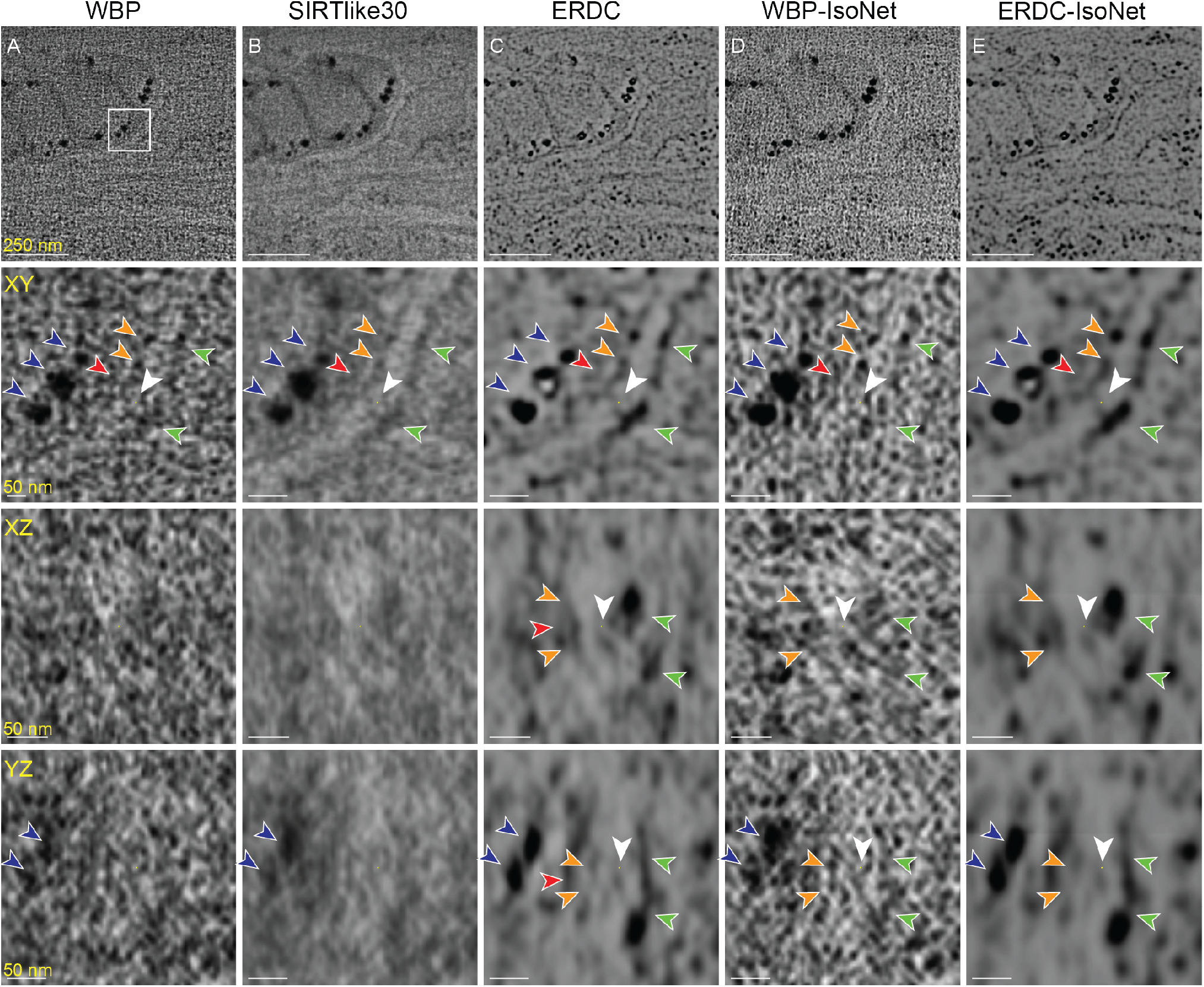
Comparing and combining deconvolution with IsoNet: Comparing (A) WBP, (B) Sirt-Like30, (C) ERDC, (D) IsoNet applied to WBP, and (E) IsoNet applied to ERDC tomogram. Top row shows the mitochondria-ER contact site. A-C are the same as in Fig 2,*D-E*. Scale bars are 250 nm. Second, third, and fourth rows show the XY, XZ, and YZ plane, respectively, at the protein tether (white arrowheads) in this contact site. Scale bars are 50 nm.

We applied IsoNet to the WBP (Fig 6*D*) and to the deconvolved dual-axis tomograms (Fig 6*E*). In the former, somewhat stronger contrast was seen for the granules in XZ and YZ views, as expected, but the images were still dominated by “salt and pepper” noise Fig 6*D*. This noise was strongly suppressed by ERDC, after which IsoNet had little further effect. If anything, there seems to be a very minor smoothing of the features with IsoNet, compared to ERDC Fig 6*E*).

From the orthoslices, the contrast from the deconvolved, dual-axis reconstructions appears to be much more isotropic in comparison with the single axis or the simple back-projections. This is additionally highlighted when looking at the corresponding power spectra (Fig 7). The improvement by combining the two axes becomes evident in the XZ and YZ planes in reciprocal space (kxkz and kykz planes, respectively) of the WBP tomogram, where the second axis fills in the missing wedge of the first (green lines in Fig 7*A*). The gaps between the sharp lines (red arrowheads, with small missing wedges in between) are filled in after ERDC (Fig 7*B*). This interpolation expresses the suppression of ghost streaks in back-projection. Only a minor effect was seen in the power spectrum after IsoNet processing (Fig 7*C,D*). IsoNet after ERDC appeared as a very mild low pass filter, which is consistent with the smoothing noted in the real space images.

**Fig 7:**
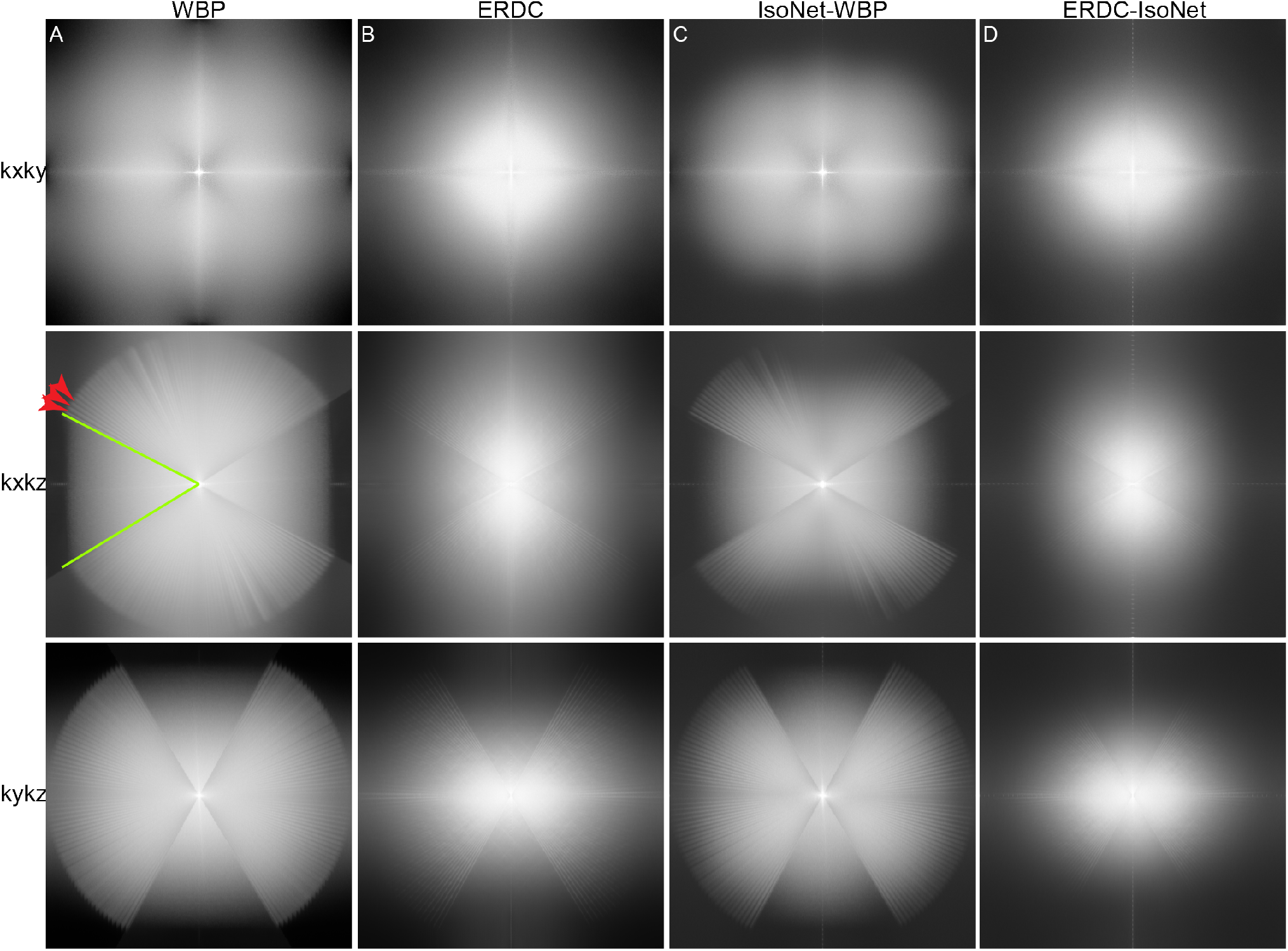
Powerspectra of the tomograms. Kxky, kxkz and kxky powerspectra of the dual-axis (A) WBP, (B) ERDC, (C) IsoNet-applied to the WBP, and (D) IsoNet applied to the ERDC tomograms. Green lines indicate the missing pyramid and red arrowheads show the small missing wedges.

These results show that ERDC is able to compensate for projection information that is still missing in dual-axis CSTET data. Clearly, the combination of dual-axis CSTET combined with deconvolution is able to reveal fine details in an 850 nm thick sample, which is otherwise inaccessible in CSTET data treated conventionally.

### 3.4 Combining cryo-SRRF with dual-axis CSTET

Finally, we correlated cryo-SRRF with dual-axis CSTET. We overlaid the FMD and the cryo-SRRF data on the medium resolution STEM map and a minimum intensity projection of the tomogram Fig 8. Cryo-SRRF provided a finer localization of the fluorescent signal in the CSTET tomogram compared to FMD. This is exemplified by the SPY650-tubulin stain, where in the FMD data, a signal more than 500 nm wide is seen, while the line width in cryo-SRRF data is close to 100 nm.

**Fig 8:**
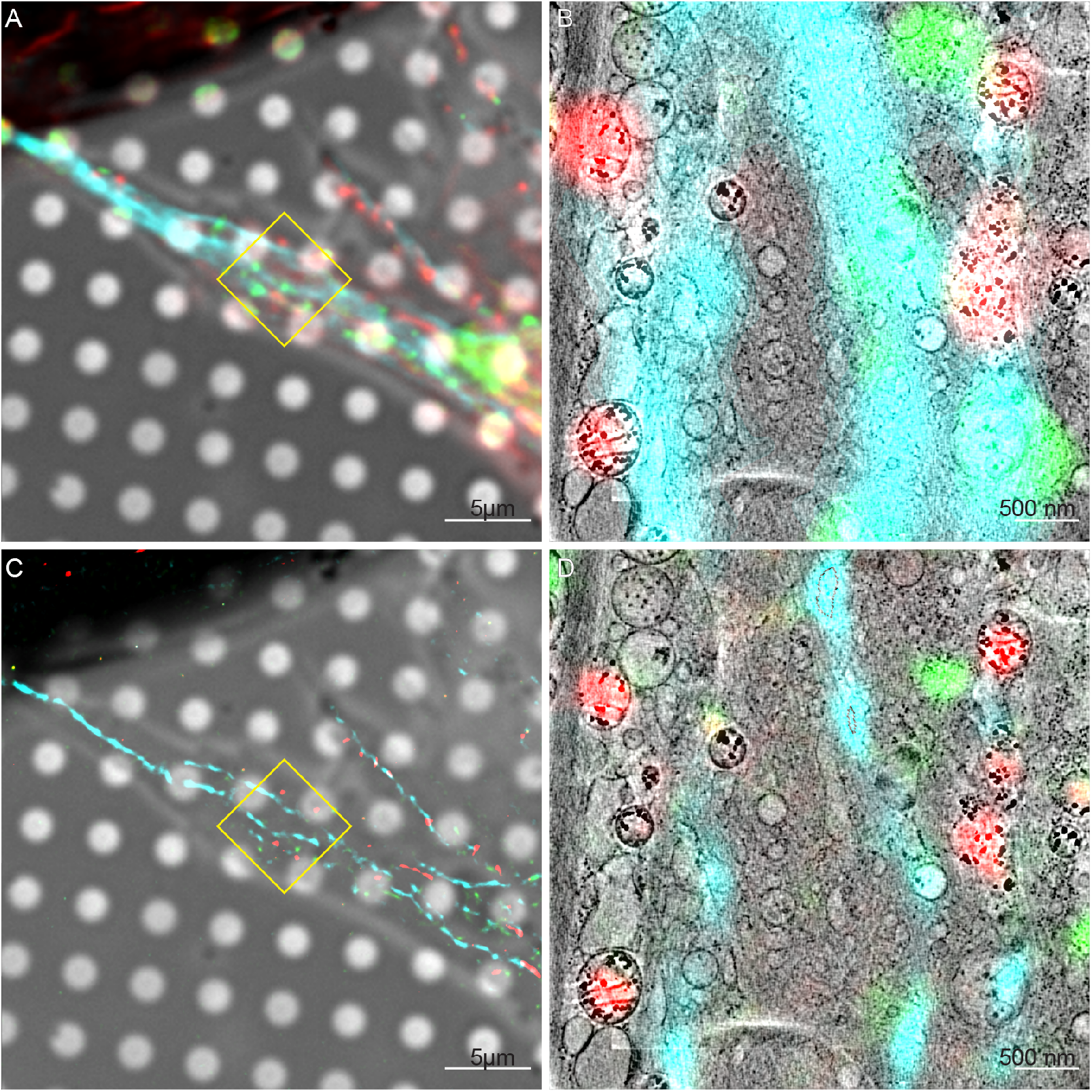
FMD and cryo-SRRF correlation with dual-axis CSTET tomogram 1. Overlay of the (A,B) deconvolved and (C,D) cryo-SRRF on the (A,C) medium resolution map and (B,D) tomogram. Red, green, and cyan mark the mitochondria, Golgi- and Golgi-derived vesicles, and microtubules, respectively. Scale bars are (A,C) 5 *μm* and (B,D) 500 nm, respectively.

## 4 Discussion

Cellular cryo-tomography faces a number of challenges that so far limit its applicability to thin extremities or lamellae. These include localization of the relevant region of interest at sufficient precision to identify macro-molecular or organelle features, severe limits on sample thickness in cryo-ET, small FOV, and the noisy nature of cryo-tomograms usually attributed to the very low permissible exposure. This work demonstrates a workflow that addresses these issues for cryo-STEM tomography.

Feature localization depends on CLEM, yet the approach suffers from the well-known resolution gap where the image of smallest resolvable feature in light microscopy may be close in size to the entire FOV in EM. In practice, the Abbe limit of 250 nm is rarely achieved in micron-thick specimens. Various high- to super-resolution methods under cryo-conditions have been tested for CLEM, such as photoactivated localization microscopy (PALM, [34]) and more generally single molecule localization microscopy (SMLM, [37]), structured illuminated microscopy (SIM, [7, 38]), spinning-disk confocal imaging [35], or super-resolution confocal imaging [39]. These methods rely on special optical hardware, fluorophores, particular buffers, and/or high laser powers for illumination. Cryogenic super-resolution optical fluctuation imaging (SOFI, [36]) avoids these problems by taking advantage of spontaneous fluorescence intensity fluctuations and their correlations. SRRF [40] combines a spatial analysis similar to STORM with a temporal analysis similar to SOFI. Both SOFI and SRRF can access the whole range of fluorescent markers, rather than those which can undergo complete blinking or survive intense laser fluxes. For cryo-microscopy applications, the modest illumination intensity relaxes the overriding concern that warming of the grid will induce sample devitrification.

In this study, we have shown that multi-color cryo-SRRF offers great promises for the localization of diverse objects of interest. We demonstrate that cryo-SRRF can reach resolutions on the order of 100-200 nm for four different fluorescence channels. Additionally, conventional deconvolution of a Z-stack acquisition provides a great improvement over simple wide-field fluorescence in the cellular cryo-CLEM context, but its resolution remains limited. To determine the resolution, we applied the Fourier Ring Correlation (FRC) value [66–68] and the edge response [52]. (The former method was also used to optimize settings during the SRRF analysis.) We note that both methods have limitations. The formal FRC value provides an unreasonably favorable estimation (especially in the case of the TMRE and BODIPY signal), while the edge response can be read as a lower bound because the distribution of fluorophores has a real width.

Additional factors should be considered together with the localization precision. For example, small lateral shifts were observed between the green and red color channels in the fluorescence deconvolution and the SRRF, when overlaid on the high resolution tomogram (Fig 8*C,D*). Such discrepancies may escape observation when overlaid on the medium resolution STEM image (Fig 8*A,B*). Lateral shifts might be introduced in deconvolution by optical refraction through the curved sample surface, essentially a local distortion of the point spread function, or in SRRF by the preferential detection of fluctuations in areas of relatively low fluorophore density.

Limitations on cell thickness and FOV are addressed by the use of STEM imaging rather than conventional wide-field TEM. STEM has the advantage of insensitivity to chromatic aberration, which obviates the need for an energy filter. Contrast is based on scattering rather than phase, which is also easier to interpret for thick specimens. A remaining limitation is the missing wedge of information, both the large missing wedge due to a limited tilt range and the small missing wedges inherent in discrete tilts. The use of a dual-axis acquisition scheme reduces the large missing wedge to a missing pyramid.

Dual-axis STEM tomograms provide significantly more detail than single-axis. It was already noted that filamentous objects lying perpendicular to the tilt axis are not well represented in single-axis tomograms, but dual-axis tomography is able to recover this information [18]. A similar situation in presented here, where only the dual-axis tomogram shows a microtubule running perpendicular to the tilt axis (Fig 5).

The conventional weighted back-projection (as well as SIRT-like filters on back-projection) unavoidably back-projects intensities throughout the reconstructed volume. With a sufficient number of projections, this spurious contrast may be insignificant, but that is not the case when their number is limited and especially when the specimen is thick. The result is that a bright point is represented as a radial array of spines, one per tilt. There is a maximum at the vertex where they meet, but in other planes, the combination of spines originating from distant densities results in a “salt and pepper” noise that appears random but is actually structural as can be seen in the plane perpendicular to the tilt axis. Additional artifacts originate from the missing wedge/pyramid. Accounting or correcting for the missing wedge is usually done by averaging of subtomogram, either of small subtomograms of individual proteins or, as in the case of IsoNet, of small cubes within the tomogram. Here we have extended the deconvolution approach to dual-axis STEM and introduced a script-based protocol to run most of the necessary steps in an automated fashion. The “salt and pepper” noise and the missing wedges are addressed by 3D deconvolution using a synthetic point spread function in 3D. Deconvolution combined with dual-axis tomography results in visibly isotropic resolution.

The results of dual-axis ERDC are very satisfactory, showing an improvement in the visibility of both high and low contrast features that were otherwise buried under noise. Moreover, the classic Z-axis elongation artifact associated with the large missing wedge is strongly suppressed. The improvement allows for the identification of molecular contact regions between the mitochondria and ER [65], even in the context of a sample almost one micron thick.

We attempted to make a further improvement in suppressing the missing wedge effects using IsoNet, which takes a complementary approach based on deep learning. Direct application of IsoNet to the WBP reconstruction was less successful than published applications to TEM tomograms. Perhaps this was due to the more severe data limitations for thick samples rather than the differences between TEM and STEM. Application of IsoNet after ERDC yielded little visible improvement in the images examined, except for mild smoothing. It is worth noting that the contrast between regions within and outside of the (single-axis) missing wedge was reduced by the IsoNet processing, suggesting that it did make some improvement toward isotropic resolution. This is highlighted by an azimuthal average of the power spectra (Fig S8). More work will be needed to establish this and to optimize parameters for practical application.

In summary, dual-axis CSTET with deconvolution and cryo-SRRF CLEM becomes a useful tool for the study of intact vitrified cells and organelles, at least up to one micron thickness. In practice, we find that the specimen thickness is limited as strictly by the vitrification as by the imaging. This raises intriguing possibilities for the study of thick sections of cells or even tissues prepared by high-pressure freezing and focused ion beam milling, as well as for correlation with 3D FIB-SEM imaging or soft X-ray tomography.

## A Supporting Material

The WBP and ERDC dual-axis tomograms are available under the EMDB deposition ID EMD-16178 and EMD-16179, respectively. The ERDC Python script and the FIJI macro for performing SRRF and calculating the FRC may be found on github: https://github.com/PKirchweger/CSTETDeconvolution and https://github.com/PKirchweger/SRRF-macro.

## B Acknowledgements

We thank Prof. Deborah Fass for continuous support throughout the project and discussions on the manuscript, Dr. Guenter Resch for discussions on the dual-axis data collection, and Prof. John Sedat for comments on the manuscript. P.K. was funded by the Austrian Science Fund (FWF) through a Schrödinger Fellowship J4449-B. For the purpose of open access, the authors have applied a CC-BY public copyright license to any Author Accepted Manuscript version arising from this submission. P.P.S. thanks the Feinberg Graduate School (FGS) at the Weizmann Institute of Science for approving the thesis internship. P.P.S. was funded in part by the Department of Science and Technology (DST), India through an INSPIRE fellowship CBS/P015927. M.E. and S.G.W. acknowledge funding from the Israel Science Foundation, grant no.1696/18, and the European Union Horizon 2020 Twinning project, IMpaCT (grant no.857203). Funding from the ERC project CryoSTEM (grant no. 101055413) is also acknowledged. M.E. is the incumbent of the Sam and Ayala Zacks Professorial Chair and head of the Irving and Cherna Moskowitz Center for Nano and Bio-Nano Imaging. The laboratory of M.E. has benefited from the historical generosity of the Harold Perlman family. Funded by the European Union. Views and opinions expressed are however those of the author(s) only and do not necessarily reflect those of the European Union or the European Research Council Executive Agency. Neither the European Union nor the granting authority can be held responsible for them.

## C Author contributions

PK: Conceptualization, Data curation, Formal Analysis, Funding acquisition, Investigation, Methodology, Software, Validation, Visualization, Writing – original draft, Writing – review & editing DM: Conceptualization, Data curation, Formal Analysis, Investigation, Methodology, Validation, Visualization, Writing – review & editing PPS: Formal Analysis, Investigation, Methodology, Software, Validation, Writing – review & editing SGW: Funding acquisition, Methodology, Supervision, Validation, Writing – review & editing ME: Conceptualization, Formal Analysis, Funding acquisition, Investigation, Methodology, Project administration, Resources, Software, Supervision, Validation, Visualization, Writing – original draft, Writing – review & editing

## References

[1] Joachim Frank. Introduction: Principles of Electron Tomography. In Joachim Frank, editor, Electron Tomography, pages 1–15. Springer New York, New York, NY, 2006. ISBN 978-0-387-69008-7. doi: 10.1007/978-0-387-69008-7{\}1. URL http://link.springer.com/10.1007/978-0-387-69008-7_1.

[2] Sharon Grayer Wolf, Lothar Houben, and Michael Elbaum. Cryo-scanning transmission electron tomography of vitrified cells. Nature Methods, 11(4):423–428, 2014. doi: 10.1038/NMETH.2842.

[3] Gerd Schneider, Peter Guttmann, Stefan Heim, Stefan Rehbein, Florian Mueller, Kunio Nagashima, J Bernard Heymann, Waltraud G Müller, and James G McNally. Three-dimensional cellular ultrastructure resolved by X-ray microscopy. Nature Methods, 7(12):985–987, 12 2010. ISSN 1548-7091. doi: 10.1038/nmeth.1533. URLhttp://www.nature.com/articles/nmeth.1533.

[4] Andreas Schertel, Nicolas Snaidero, Hong-Mei Han, Torben Ruhwedel, Michael Laue, Markus Grabenbauer, and Wiebke Möbius. Cryo FIB-SEM: Volume imaging of cellular ultrastructure in native frozen specimens. Journal of Structural Biology, 184(2):355–360, 11 2013. ISSN 10478477. doi: 10.1016/j.jsb.2013.09.024. URL https://linkinghub.elsevier.com/retrieve/pii/S1047847713002657.

[5] Sanja Sviben, Assaf Gal, Matthew A. Hood, Luca Bertinetti, Yael Politi, Mathieu Bennet, Praveen Krishnamoorthy, Andreas Schertel, Richard Wirth, Andrea Sorrentino, Eva Pereiro, Damien Faivre, and André Scheffel. A vacuole-like compartment concentrates a disordered calcium phase in a key coccolithophorid alga. Nature Communications, 7(1):11228, 9 2016. ISSN 2041-1723. doi: 10.1038/ncomms11228.

[6] Neta Varsano, Keren Kahil, Heden Haimov, Katya Rechav, Lia Addadi, and Steve Weiner. Characterization of the growth plate-bone interphase region using cryo-FIB SEM 3D volume imaging. Journal of Structural Biology, 213(4):107781, 12 2021. ISSN 10478477. doi: 10.1016/j.jsb.2021.107781.

[7] David P. Hoffman, Gleb Shtengel, C. Shan Xu, Kirby R. Campbell, Melanie Freeman, Lei Wang, Daniel E. Milkie, H. Amalia Pasolli, Nirmala Iyer, John A. Bogovic, Daniel R. Stabley, Abbas Shirinifard, Song Pang, David Peale, Kathy Schaefer, Wim Pomp, Chi-Lun Chang, Jennifer Lippincott-Schwartz, Tom Kirchhausen, David J. Solecki, Eric Betzig, and Harald F. Hess. Correlative three-dimensional super-resolution and block-face electron microscopy of whole vitreously frozen cells. Science, 367(6475), 1 2020. ISSN 0036-8075. doi:10.1126/science.aaz5357. URL https://www.science.org/doi/10.1126/science.aaz5357.

[8] Michael Elbaum. Expanding horizons of cryo-tomography to larger volumes. Current Opinion in Microbiology, 43:155–161, 6 2018. ISSN 13695274. doi: 10.1016/j.mib.2018.01.001. URL https://linkinghub.elsevier.com/retrieve/pii/S1369527417300498.

[9] Neta Varsano and Sharon Grayer Wolf. Electron microscopy of cellular ultrastructure in three dimensions. Current Opinion in Structural Biology, 76:102444, 10 2022. ISSN 0959440X. doi: 10.1016/j.sbi.2022.102444.

[10] Felix J.B. Bäuerlein and Wolfgang Baumeister. Towards Visual Proteomics at High Resolution. Journal of Molecular Biology, 433(20):167187, 10 2021. ISSN 0022-2836. doi: 10.1016/J.JMB.2021.167187.

[11] David Kirchenbuechler, Yael Mutsafi, Ben Horowitz, Smadar Levin-Zaidman, Deborah Fass, Sharon G. Wolf, and Michael Elbaum. Cryo-STEM Tomography of Intact Vitrified Fibroblasts. AIMS Biophysics, 2(3):259–273, 2015. ISSN 2377-9098. doi: 10.3934/biophy.2015.3.259. URL http://www.aimspress.com/article/10.3934/biophy.2015.3.259.

[12] Sharon Grayer Wolf, Eyal Shimoni, Michael Elbaum, and Lothar Houben. STEM Tomography in Biology. In Eric Hanssen, editor, Cellular Imaging: Electron Tomography and Related Techniques, pages 33–60. Springer International Publishing, 2018. ISBN 978-3-319-68997-5. doi: 10.1007/978-3-319-68997-5{\}2. URL http://link.springer.com/10.1007/978-3-319-68997-5_2 https://doi.org/10.1007/978-3-319-68997-5_2.

[13] Michael Elbaum, Shahar Seifer, Lothar Houben, Sharon G Wolf, and Peter Rez. Toward Compositional Contrast by Cryo-STEM. Accounts of Chemical Research, 9 2021. ISSN 0001-4842. doi: 10.1021/acs.accounts.1c00279. URLhttps://doi.org/10.1021/acs.accounts.1c00279.

[14] Emelie Fogelqvist, Mikael Kördel, Valentina Carannante, Björn Önfelt, and Hans M. Hertz. Laboratory cryo x-ray microscopy for 3D cell imaging. Scientific Reports, 7(1):13433, 12 2017. ISSN 2045-2322. doi: 10.1038/s41598-017-13538-2.

[15] Johann Radon. Ü ber die Bestimmung von Funktionen durch ihre Integralwerte längs gewisser Mannigfaltigkeiten. Berichte senschaften, 69:262–277, 1917. über die Verhandlungen der Sächsische Akademie der Wis-

[16] Pawel A. Penczek. Fundamentals of Three-Dimensional Reconstruction from Projections. In Methods in Enzymology, volume 482, pages 1–33. Academic Press Inc., 2010. doi: 10.1016/S0076-6879(10)82001-4. URLhttps://linkinghub.elsevier.com/retrieve/pii/S0076687910820014.

[17] Pawel Penczek, Michael Marko, Karolyn Buttle, and Joachim Frank. Double-tilt electron tomography. Ultramicroscopy, 60(3):393–410, 1995.

[18] David N. Mastronarde. Dual-axis tomography: An approach with alignment methods that preserve resolution. Journal of Structural Biology, 120(3):343–352, 1997.

[19] Sébastien Phan, Daniela Boassa, Phuong Nguyen, Xiaohua Wan, Jason Lanman, Albert Lawrence, and Mark H Ellisman. 3D reconstruction of biological structures: automated procedures for alignment and reconstruction of multiple tilt series in electron tomography. Advanced Structural and Chemical Imaging, 2(1):8, 12 2016. ISSN 2198-0926. doi: 10.1186/s40679-016-0021-2. URL https://link.springer.com/10.1186/s40679-016-0021-2.

[20] Alioscka A. Sousa, Afrouz A. Azari, Guofeng Zhang, and Richard D. Leapman. Dual-axis electron tomography of biological specimens: Extending the limits of specimen thickness with bright-field STEM imaging. Journal of Structural Biology, 174(1):107–114, 2011. ISSN 10478477. doi: 10.1016/j.jsb.2010.10.017. URLhttp://dx.doi.org/10.1016/j.jsb.2010.10.017.

[21] Reinhard Rachel, Paul Walther, Christine Maaßen, Ingo Daberkow, Masahiro Matsuoka, and Ralph Witzgall. Dual-axis STEM tomography at 200 kV: Setup, performance, limitations. Journal of Structural Biology, 211(3):107551, 2020. ISSN 10958657. doi: 10.1016/j.jsb.2020.107551. URL https://doi.org/10.1016/j.jsb.2020.107551.

[22] Sophie L. Winter and Petr Chlanda. Dual-axis Volta phase plate cryo-electron tomography of Ebola virus-like particles reveals actin-VP40 interactions. Journal of Structural Biology, 213(2):107742, 6 2021. ISSN 10958657. doi: 10.1016/j.jsb.2021.107742.

[23] Michael Elbaum. Quantitative Cryo-Scanning Transmission Electron Microscopy of Biological Materials. Advanced Materials, 30(41):1706681, 10 2018. ISSN 09359648. doi: 10.1002/adma.201706681.

[24] Murat Nulati Yesibolati, Simone Laganá, Shima Kadkhodazadeh, Esben Kirk Mikkelsen, Hongyu Sun, Takeshi Kasama, Ole Hansen, Nestor J. Zaluzec, and Kristian Mølhave. Electron inelastic mean free path in water. Nanoscale, 12(40):20649–20657, 2020. ISSN 2040-3364. doi: 10.1039/D0NR04352D.

[25] Michael Marko, Chyongere Hsieh, Richard Schalek, Joachim Frank, and Carmen Mannella. Focused-ion-beam thinning of frozen-hydrated biological specimens for cryo-electron microscopy. Nature Methods, 4(3):215–217, 3 2007. ISSN 1548-7091. doi: 10.1038/nmeth1014. URL http://www.nature.com/articles/nmeth1014.

[26] Elizabeth Villa, Miroslava Schaffer, Jürgen M. Plitzko, and Wolfgang Baumeister. Opening windows into the cell: focused-ion-beam milling for cryo-electron tomography. Current Opinion in Structural Biology, 23(5):771–777, 10 2013. ISSN 0959-440X. doi: 10.1016/J.SBI.2013.08.006.

[27] Julia Mahamid, Stefan Pfeffer, Miroslava Schaffer, Elizabeth Villa, Radostin Danev, Luis Kuhn Cuellar, Friedrich Förster, Anthony A. Hyman, Jürgen M. Plitzko, and Wolfgang Baumeister. Visualizing the molecular sociology at the HeLa cell nuclear periphery. Science, 351(6276):969–972, 2 2016. ISSN 0036-8075. doi: 10.1126/science.aad8857.

[28] Sharon Grayer Wolf, Yael Mutsafi, Tali Dadosh, Tal Ilani, Zipora Lansky, Ben Horowitz, Sarah Rubin, Michael Elbaum, and Deborah Fass. 3D visualization of mitochondrial solidphase calcium stores in whole cells. eLife, 6, 11 2017. ISSN 2050-084X. doi: 10.7554/eLife.29929. URL https://elifesciences.org/articles/29929.

[29] Barnali Waugh, Sharon G. Wolf, Deborah Fass, Eric Branlund, Zvi Kam, John W. Sedat, and Michael Elbaum. Three-dimensional deconvolution processing for STEM cryotomography. Proceedings of the National Academy of Sciences, 117(44):27374–27380, 11 2020. ISSN 0027-8424. doi: 10.1073/PNAS.2000700117. URLhttps://www.pnas.org/content/117/44/27374.

[30] M. Arigovindan, J. C. Fung, D. Elnatan, V. Mennella, Y.-H. M. Chan, M. Pollard, E. Branlund, J. W. Sedat, and D. A. Agard. High-resolution restoration of 3D structures from widefield images with extreme low signal-to-noise-ratio. Proceedings of the National Academy of Sciences, 110(43):17344–17349, 10 2013. ISSN 0027-8424. doi: 10.1073/pnas.1315675110. URL http://www.pnas.org/cgi/doi/10.1073/pnas.1315675110.

[31] Matthew Croxford, Michael Elbaum, Muthuvel Arigovindan, Zvi Kam, David Agard, Elizabeth Villa, and John Sedat. Entropy-regularized deconvolution of cellular cryotransmission electron tomograms. Proceedings of the National Academy of Sciences, 118(50): e2108738118, 12 2021. ISSN 0027-8424. doi: 10.1073/pnas.2108738118. URLhttp://www.pnas.org/lookup/doi/10.1073/pnas.2108738118.

[32] Cindi L. Schwartz, Vasily I. Sarbash, Fazoil I. Ataullakhanoc, J. Richard Mcintosh, and Daniela Nicastro. Cryo-fluorescence microscopy facilitates correlations between light and cryo-electron microscopy and reduces the rate of photobleaching. Journal of Microscopy, 227 (2):98–109, 8 2007. ISSN 0022-2720. doi: 10.1111/j.1365-2818.2007.01794.x. URLhttps://onlinelibrary.wiley.com/doi/10.1111/j.1365-2818.2007.01794.x.

[33] Peijun Zhang. Correlative cryo-electron tomography and optical microscopy of cells. Current Opinion in Structural Biology, 23(5):763–770, 10 2013. ISSN 0959440X. doi: 10.1016/j.sbi.2013.07.017. URLhttps://linkinghub.elsevier.com/retrieve/pii/S0959440X13001504.

[34] Yi-Wei Chang, Songye Chen, Elitza I Tocheva, Anke Treuner-Lange, Stephanie Löbach, Lotte Søgaard-Andersen, and Grant J Jensen. Correlated cryogenic photoactivated localization microscopy and cryo-electron tomography. Nature Methods, 11(7):737–739, 7 2014. ISSN 1548-7091. doi: 10.1038/nmeth.2961. URLhttp://www.nature.com/articles/nmeth.2961.

[35] Jan Arnold, Julia Mahamid, Vladan Lucic, Alex de Marco, Jose-Jesus Fernandez, Tim Laugks, Tobias Mayer, Anthony A. Hyman, Wolfgang Baumeister, and Jürgen M. Plitzko. Site-Specific Cryo-focused Ion Beam Sample Preparation Guided by 3D Correlative Microscopy. Biophysical Journal, 110(4):860–869, 2 2016. ISSN 00063495. doi: 10.1016/j.bpj.2015.10.053. URLhttps://linkinghub.elsevier.com/retrieve/pii/S0006349515011637.

[36] Felipe Moser, Vojtech Prazák, Valerie Mordhorst, Débora M. Andrade, Lindsay A. Baker, Christoph Hagen, Kay Grünewald, and Rainer Kaufmann. Cryo-SOFI enabling low-dose super-resolution correlative light and electron cryo-microscopy. Proceedings of the National Academy of Sciences of the United States of America, 116(11):4804–4809, 2019. ISSN 10916490. doi: 10.1073/pnas.1810690116.

[37] Maarten W. Tuijtel, Abraham J. Koster, Stefan Jakobs, Frank G.A. Faas, and Thomas H. Sharp. Correlative cryo super-resolution light and electron microscopy on mammalian cells using fluorescent proteins. Scientific Reports, 9(1), 2019. ISSN 20452322. doi: 10.1038/s41598-018-37728-8.

[38] Michael A. Phillips, Maria Harkiolaki, David Miguel Susano Pinto, Richard M. Parton, Ana Palanca, Manuel Garcia-Moreno, Ilias Kounatidis, John W. Sedat, David I. Stuart, Alfredo Castello, Martin J. Booth, Ilan Davis, and Ian M. Dobbie. CryoSIM: super-resolution 3D structured illumination cryogenic fluorescence microscopy for correlated ultrastructural imaging. Optica, 7(7):802, 7 2020. ISSN 2334-2536. doi: 10.1364/OPTICA.393203. URL https://www.osapublishing.org/abstract.cfm?URI=optica-7-7-802.

[39] Danielle L. Sexton, Steffen Burgold, Andreas Schertel, and Elitza I. Tocheva. Superresolution confocal cryo-CLEM with cryo-FIB milling for in situ imaging of Deinococcus radiodurans. Current Research in Structural Biology, 4:1–9, 1 2022. ISSN 2665928X. doi: 10.1016/j.crstbi.2021.12.001. URLhttps://linkinghub.elsevier.com/retrieve/pii/S2665928X21000295.

[40] Nils Gustafsson, Siân Culley, George Ashdown, Dylan M. Owen, Pedro Matos Pereira, and Ricardo Henriques. Fast live-cell conventional fluorophore nanoscopy with ImageJ through super-resolution radial fluctuations. Nature Communications, 7:1–9, 2016. ISSN 20411723. doi: 10.1038/ncomms12471.

[41] Siân Culley, Kalina L. Tosheva, Pedro Matos Pereira, and Ricardo Henriques. SRRF: Universal live-cell super-resolution microscopy. International Journal of Biochemistry and Cell Biology, 101(March):74–79, 2018. ISSN 18785875. doi: 10.1016/j.biocel.2018.05.014. URL https://doi.org/10.1016/j.biocel.2018.05.014.

[42] Zipora Lansky, Yael Mutsafi, Lothar Houben, Tal Ilani, Gad Armony, Sharon G. Wolf, and Deborah Fass. 3D mapping of native extracellular matrix reveals cellular responses to the microenvironment. Journal of Structural Biology: X, 1:100002, 1 2019. ISSN 2590-1524. doi: 10.1016/J.YJSBX.2018.100002.

[43] Florian Fässler, Bettina Zens, Robert Hauschild, and Florian K.M. Schur. 3D printed cell culture grid holders for improved cellular specimen preparation in cryo-electron microscopy. Journal of Structural Biology, 212(3):107633, 12 2020. ISSN 1047-8477. doi: 10.1016/J.JSB.2020.107633.

[44] Martin Schorb, Leander Gaechter, Ori Avinoam, Frank Sieckmann, Mairi Clarke, Cecilia Bebeacua, Yury S. Bykov, Andreas F.P. Sonnen, Reinhard Lihl, and John A.G. Briggs. New hardware and workflows for semi-automated correlative cryo-fluorescence and cryo-electron microscopy/tomography. Journal of Structural Biology, 197(2):83–93, 2 2017. ISSN 1047-8477. doi: 10.1016/J.JSB.2016.06.020.

[45] Johannes Schindelin, Ignacio Arganda-Carreras, Erwin Frise, Verena Kaynig, Mark Longair, Tobias Pietzsch, Stephan Preibisch, Curtis Rueden, Stephan Saalfeld, Benjamin Schmid, Jean-Yves Tinevez, Daniel James White, Volker Hartenstein, Kevin Eliceiri, Pavel Tomancak, and Albert Cardona. Fiji: an open-source platform for biological-image analysis. Nature Methods, 9(7):676–682, 7 2012. ISSN 1548-7091. doi: 10.1038/nmeth.2019. URL http://www.nature.com/articles/nmeth.2019.

[46] Kang Li. The image stabilizer plugin for ImageJ, 2 2008. URLhttp://www.cs.cmu.edu/~kangli/code/Image_Stabilizer.html.

[47] H. Kirshner, F. Aguet, D. Sage, and M. Unser. 3-D PSF fitting for fluorescence microscopy: Implementation and localization application. Journal of Microscopy, 249(1):13–25, 1 2013. ISSN 00222720. doi: 10.1111/j.1365-2818.2012.03675.x. URL http://www.ncbi.nlm.nih.gov/pubmed/23126323https://onlinelibrary.wiley.com/doi/10.1111/j.1365-2818.2012.03675.x.

[48] Daniel Sage, Lauréne Donati, Ferréol Soulez, Denis Fortun, Guillaume Schmit, Arne Seitz, Romain Guiet, Cédric Vonesch, and Michael Unser. DeconvolutionLab2: An open-source software for deconvolution microscopy. Methods, 115:28–41, 2 2017. ISSN 1046-2023. doi: 10.1016/J.YMETH.2016.12.015.

[49] Arthur Edelstein, Nenad Amodaj, Karl Hoover, Ron Vale, and Nico Stuurman. Computer Control of Microscopes Using μManager. Current Protocols in Molecular Biology, 92(1), 10 2010. ISSN 1934-3639. doi: 10.1002/0471142727.mb1420s92.

[50] Arthur D Edelstein, Mark A Tsuchida, Nenad Amodaj, Henry Pinkard, Ronald D Vale, and Nico Stuurman. Advanced methods of microscope control using μManager software. Journal of Biological Methods, 1(2):e10, 11 2014. ISSN 2326-9901. doi: 10.14440/jbm.2014.36. URL https://jbmethods.org/jbm/article/view/36.

[51] Romain F. Laine, Kalina L. Tosheva, Nils Gustafsson, Robert D.M. Gray, Pedro Almada, David Albrecht, Gabriel T. Risa, Fredrik Hurtig, Ann Christin Lindås, Buzz Baum, Jason Mercer, Christophe Leterrier, Pedro M. Pereira, Siân Culley, and Ricardo Henriques. NanoJ: A high-performance open-source super-resolution microscopy toolbox. Journal of Physics D: Applied Physics, 52(16), 2019. ISSN 13616463. doi: 10.1088/1361-6463/ab0261.

[52] Steven W. Smith. Special Imaging Techniques. In Digital Signal Processing, pages 423–450. Elsevier, 2003. doi: 10.1016/B978-0-7506-7444-7/50062-5.

[53] David N. Mastronarde. SerialEM: A program for automated tilt series acquisition on Tecnai microscopes using prediction of specimen position. Microscopy and Microanalysis, 9(SUPPL. 2):1182–1183, 2003. doi: 10.1017/S1431927603445911.

[54] David N. Mastronarde. Automated electron microscope tomography using robust prediction of specimen movements. Journal of Structural Biology, 152(1):36–51, 10 2005. ISSN 1047-8477. doi: 10.1016/J.JSB.2005.07.007.

[55] Wim J.H. Hagen, William Wan, and John A.G. Briggs. Implementation of a cryo-electron tomography tilt-scheme optimized for high resolution subtomogram averaging. Journal of Structural Biology, 197(2):191–198, 2 2017. ISSN 1047-8477. doi: 10.1016/J.JSB.2016.06.007.

[56] James R. Kremer, David N. Mastronarde, and J. Richard McIntosh. Computer visualization of three-dimensional image data using IMOD. Journal of Structural Biology, 116(1):71–76, 1996.

[57] David N. Mastronarde and Susannah R. Held. Automated tilt series alignment and tomographic reconstruction in IMOD. Journal of Structural Biology, 197(2):102–113, 2 2017.

[58] Gengsheng L. Zeng. A filtered backprojection algorithm with characteristics of the iterative landweber algorithm. Medical Physics, 39(2):603–607, 2 2012. ISSN 0094-2405. doi: https://doi.org/10.1118/1.3673956. URL https://doi.org/10.1118/1.3673956.

[59] Thomas D. Goddard, Conrad C. Huang, Elaine C. Meng, Eric F. Pettersen, Gregory S. Couch, John H. Morris, and Thomas E. Ferrin. UCSF ChimeraX: Meeting modern challenges in visualization and analysis. Protein Science, 27(1):14–25, 1 2018. ISSN 09618368. doi: 10.1002/pro.3235. URL https://onlinelibrary.wiley.com/doi/10.1002/pro.3235.

[60] Eric F. Pettersen, Thomas D. Goddard, Conrad C. Huang, Elaine C. Meng, Gregory S. Couch, Tristan I. Croll, John H. Morris, and Thomas E. Ferrin. UCSF ChimeraX : Structure visualization for researchers, educators, and developers. Protein Science, 30(1):70–82, 1 2021. ISSN 0961-8368. doi: 10.1002/pro.3943. URLhttps://onlinelibrary.wiley.com/doi/10.1002/pro.3943.

[61] Hans Chen, Warren K. Clyborne, John W. Sedat, and David A. Agard. PRIISM: an integrated system for display and analysis of 3-D microscope images. In Raj S. Acharya, Carol J. Cogswell, and Dmitry B. Goldgof, editors, Biomedical Image Processing and Three-Dimensional Microscopy, volume 1660, pages 784–790, 6 1992. doi: 10.1117/12.59604. URL http://proceedings.spiedigitallibrary.org/proceeding.aspx?articleid=987191.

[62] Perrine Paul-Gilloteaux, Xavier Heiligenstein, Martin Belle, Marie-Charlotte Domart, Banafshe Larijani, Lucy Collinson, Graça Raposo, and Jean Salamero. eC-CLEM: flexible multidimensional registration software for correlative microscopies. Nature Methods, 14(2): 102–103, 2 2017. ISSN 1548-7091. doi: 10.1038/nmeth.4170. URLhttp://www.nature.com/articles/nmeth.4170.

[63] Fabrice de Chaumont, Stéphane Dallongeville, Nicolas Chenouard, Nicolas Hervé, Sorin Pop, Thomas Provoost, Vannary Meas-Yedid, Praveen Pankajakshan, Timothée Lecomte, Yoann Le Montagner, Thibault Lagache, Alexandre Dufour, and Jean-Christophe Olivo-Marin. Icy: an open bioimage informatics platform for extended reproducible research. Nature Methods, 9(7):690–696, 7 2012. ISSN 1548-7091. doi: 10.1038/nmeth.2075. URL http://www.nature.com/articles/nmeth.2075.

[64] Yun-Tao Liu, Heng Zhang, Hui Wang, Chang-Lu Tao, Guo-Qiang Bi, and Z Hong Zhou. Isotropic Reconstruction of Electron Tomograms with Deep Learning. bioRxiv, page 2021.07.17.452128, 2021. URLhttp://biorxiv.org/content/early/2021/07/19/2021.07.17.452128.abstract.

[65] Luca Scorrano, Maria Antonietta De Matteis, Scott Emr, Francesca Giordano, György Hajnóczky, Benoît Kornmann, Laura L. Lackner, Tim P. Levine, Luca Pellegrini, Karin Reinisch, Rosario Rizzuto, Thomas Simmen, Harald Stenmark, Christian Ungermann, and Maya Schuldiner. Coming together to define membrane contact sites. Nature Communications, 10(1):1287, 12 2019. ISSN 2041-1723. doi: 10.1038/s41467-019-09253-3. URL http://www.nature.com/articles/s41467-019-09253-3.

[66] W. O. Saxton and W. Baumeister. The correlation averaging of a regularly arranged bacterial cell envelope protein. Journal of Microscopy, 127(2):127–138, 8 1982. ISSN 00222720. doi: 10.1111/j.1365-2818.1982.tb00405.x.

[67] Marin Van Heel, W Keegstra, W Schutter, and E.F.J. van Bruggen. Arthropod hemocyanin structures studied by image analysis. In E.J. Wood, editor, Structure and Function of Invertebrate Respiratory Proteins, EMBO Workshop 1982, pages 69–73. 1982. ISBN 9783718601554.

[68] Alex Herbert and Oliver Burri. Fourier Ring Correlation ImageJ Plugin, 2016. URLhttps://github.com/BIOP/ijp-frc.

