## Supplemental Information for "Bridging the light-electron resolution gap with correlative cryo-SRRF and dual-axis cryo-STEM tomography"

### 1 Supplemental methods

#### 1.1 SRRF analysis

For SRRF processing, the input datasets are a sequential fast series of images of the fluorescently labelled sample. Images in the series must be background corrected, especially to remove any kind of stripes arising from the use of a sCMOS camera, and also aligned against drift of the microscope stage. The software was used as an plugin to ImageJ [2–4].

SRRF is based on the assumption that point sources when imaged are convolved with the microscope PSF, which endows them with a higher degree of radial symmetry compared to the background. It differs from other single molecule localisation approaches such as PALM or STORM in this aspect, that instead of detecting and localising a single radially symmetric point, it calculates the degree of gradient of convergence (designated as 'radiality') on a sub-pixel basis for all the pixels in the entire frame. For example, for an original frame of 3x3 pixels, and a "radiality magnification factor" of 3, we ask the algorithm to calculate a 'radiality' map on a subpixelated frame of 9x9 pixels. The determination of the magnification factor, the radius for which the radial symmetry estimation will be performed, and the number of symmetry axes to be used for calculation should be consistent with the structure under study, and these parameters need to be estimated. Similarly, use of other features provided by the software was subject to the signal to noise ratio (SNR) of the obtained data sets under study. To eliminate the chances of noise displaying radial symmetry, the temporal correlation estimation features (such as the temporal radiality pairwise product mean (TRPPM)) implemented in the plugin were used, and the number of frames that provide the best analysis was evaluated using different temporal block sizes. This optimization of parameters was done using Fourier Ring Correlation (FRC) tests [5]. Frames of the image series were divided into odd and even sequences and processed by SRRF using a range of parameters, and the FRC was used to select parameters that yield highest resolution based on the 1/7 correlation value [6]. A simple ImageJ macro developed aids this process of finding the best parameters for the structures and fluorophores under study. The macro denoises the data set, splits the series into even and odd frames, estimates drift and performs SRRF analysis using a set of parameters from both sequences, giving an FRC value. Using iterations of this macro and different choices of parameters, this FRC value has to be minimised to obtain the best resolution for a particular dataset. The set of parameters with the lowest 1/7 value was used to perform the final analysis. For all four data sets in this study, radiality magnifiction was set to 10. The final set of parameters for the SRRF analysis and their resolutions are summarized in Table S1. The macro can be downloaded here: https://github.com/PKirchweger/SRRF-macro.

Table S1: Final SRRF settings and resolution estimates

| Dataset | SRRF-settings* | | | FRC | | Edge response** [nm] | |
|---|---|---|---|---|---|---|---|
| | RR | SY | GS | 1/Nyquist | [nm] | Cryo-SRRF | FMD |
| Hoechst | 1.5 | 2 | N | 6.392 | 166 | 188 (59) | 612 (38) |
| Bodipy-Ceramide | 0.75 | 8 | Y | 2.838 | 74 | 155 (25) | 409 (53) |
| TMRE | 1 | 8 | N | 2.128 | 55 | 125 (10) | 449 (50) |
| SPY650-Tubulin | 1 | 2 | N | 5.852 | 152 | 175 (21) | 404 (82) |

*Ring Radius (RR), Axes of Symmetry (SY), Gradient Smoothing (GS)
**Standard deviation in parenthesis.

#### 1.2 Tomogram reconstruction

SIRT-like filter (equivalent to 30 iterations) was used for visualization and tomogram combination. Reconstruction settings included super-sampling by 2 for input and reconstruction and high-frequency filtering option "Hamming-like filter (as in tomo3d) starting from: 0.0". The a- and b-axis tomograms were combined using fiducial markers on one side, and fiducials with a higher residual than 8.0 were excluded. The patch sizes of 300x300x149 pixels and Kernel filtering with sigma to 2.0 were chosen. The estimated axis rotation angle for tomogram 1 and 2 were 89.94° and 89.97°, respectively. Large residual vectors with low cross-correlation values were manually deleted from the patch vector model before the Matchorwarp step in etomo.

For subsequent ERDC, the gold fiducials were deleted from the aligned tilt series using findbeads3d (IMOD), to reduce the impact of high-intensity features, and a WBP tomogram was reconstructed for both axes. Reconstruction settings included super-sampling by 2 for input and reconstruction and high-frequency filtering option "Hamming-like filter (as in tomo3d) starting from: 0.0". The tomograms were combined using the patches found from the in previous run by "Restart at Matchorwarp".

Finally, both combined tomograms were rotated around the x-axis and trimmed to the actual thickness of the cell.

#### 1.3 ERDC python script

The deconvolution script requires a pre-calculated single-probe PSF, a simulated probe which is created by the Diffraction PSF 3D plugin [7] in FIJI [2]. Settings included the numerical aperture (probe semi-convergence angle) of 0.0012, and wavelength 0.002 nm (for 300 kV). Longitudinal Spherical Aberration was left at 0 nm. Pixel size and slice spacing were set to match the reconstructed tomogram (namely 4.084 nm/pixel), and the depth was set to 1200 slices. The resulting volume was exported using the MRC-writer plugin from TomoJ [8, 9], and the pixel size of the PSF was adapted with alterheader (IMOD).

Certain settings must be set manually in the "input-values.txt", such as if a single- or dual-axis tomogram is provided, the number of iterations, the requested smoothing factors, the auxiliary smoothing factor, whether to do a summed FFT and PS, how many threads, and how much memory is available. For the single-axis tomogram, the script needs the single-probe PSF, the single-axis tomogram, and the refined tilt angles (the .tlt file from IMOD). For the dual-axis tomogram, the script uses the single-probe PSF, the dual-axis tomogram, the refined tilt angles from both axes (the .tlt files from IMOD), and the patchcorr.log (IMOD) as input. The script rotates the single-probe PSF according to the tilt angles using the rotatevol program (IMOD) and outputs a file with the size of the tomogram for each tilt-step. It then generates a multi-probe PSF of the first axis by summing the rotated PSFs using "clip add" (IMOD). In order to deconvolve dual-axis tomograms, a second multi-probe PSF is generated using the .tlt file from the b-axis and flipped using "clip flipxy" (IMOD). This second multi-probe PSF is rotated according to the refined x-axis rotation angle from the tomogram combination step by subtracting 90 minus the axis rotation angle. For a dual-axis tomogram, both multi-probe PSFs are then merged (with "clip add", IMOD) to a final PSF. Then the program FTransform3D (Priism, [10]) converts the CSTET final PSF into an optical transfer function (OTF). Finally, the core2_decon program (Priism, [9–11]) performs the actual deconvolution. Optionally, the script can calculate a summed Power Spectrum or a summed FFT of the input and output tomograms.

The script and the additional required files can be downloaded here:
https://github.com/PKirchweger/CSTET_Deconvolution.

#### 1.4 Azimuthal averages

Azimuthal averages were calculated from the power spectra using the Azimuthal Average plugin [12] in Fiji [2]. The following settings were used: the X and Y center were kept at 512 pixels, the outer radius was set to 512 pixels, the starting and integration angle were set to 0° and 180°, respectively, the number of bins was set to 360, to have a step every 1°, and spatial calibration option was chosen.

The resulting values were then overlaid and plotted with the Normalized Integrated Intensity over the Angles.

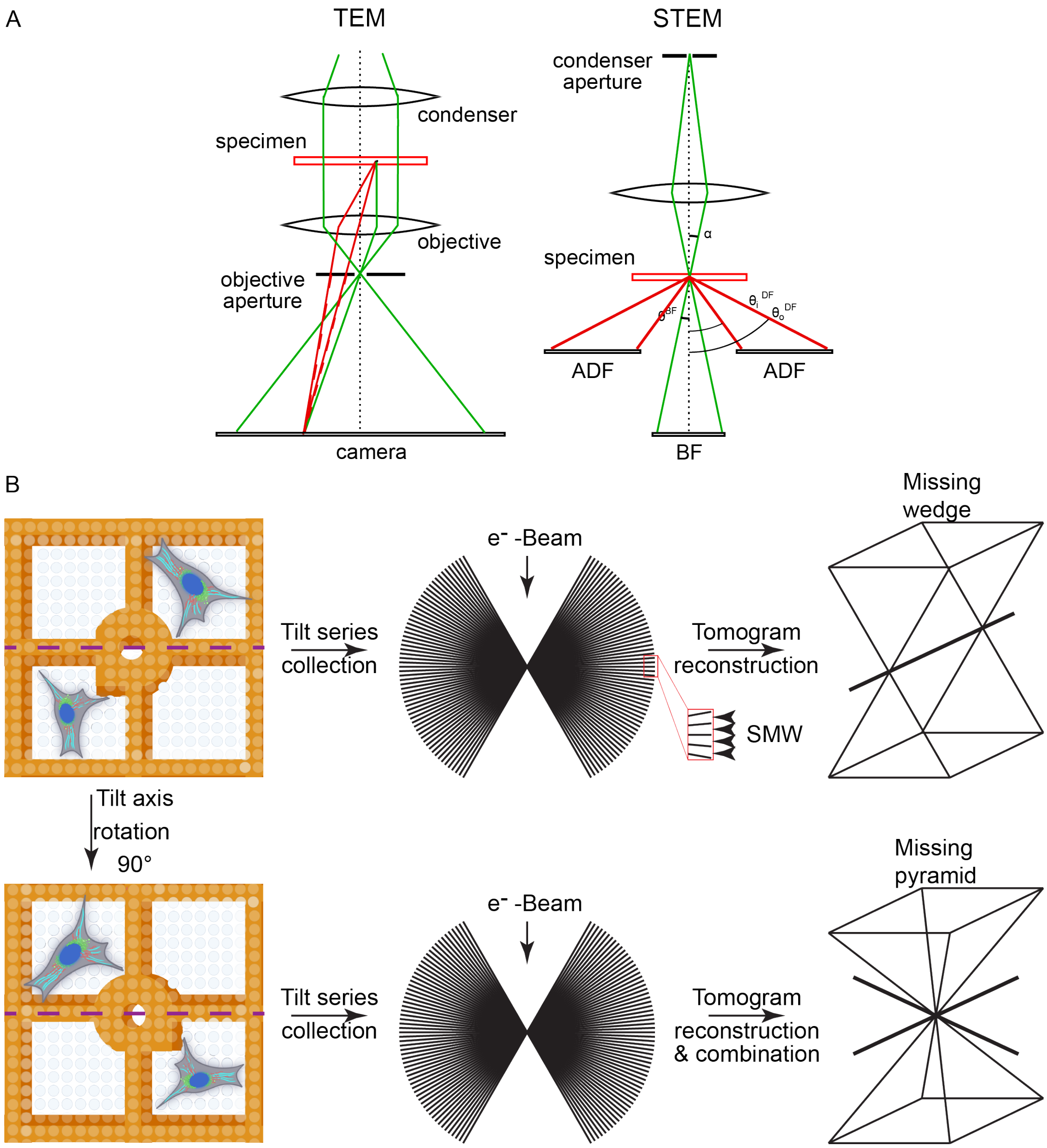


Fig S1: Comparing TEM and STEM, and schematic representation of dual-axis tomogramphy. (A) In TEM, the specimen is illuminated with a wide, near-parallel beam, while an objective lens creates the image on the camera. Phase contrast is generated by the interference between scattered (dashed red line) and unscattered (green) waves. Contrast is defocus-dependent. STEM, on the other hand, uses a focused beam that is rastered across the specimen. The scattered electrons are recorded on detectors in a pixel-by-pixel manner. Several types of detectors are positioned to collected electrons scattered to different angles. The choice of the beam and detector angle determine image properties. (Image adapted from [1]). (B) Schematic representation of dual-axis tomography. The tilt-series of the first axis is recorded and the sample is rotated 90°. Subsequently, a second tilt-series is recorded. Reconstructing the tomogram results in a so-called missing wedge of information on the top and bottom of the tomogram (in real space), and small missing wedges between the discrete tilt-steps. Reconstruction of a dual-axis tomogram results reduces the missing wedge to a missing pyramid.

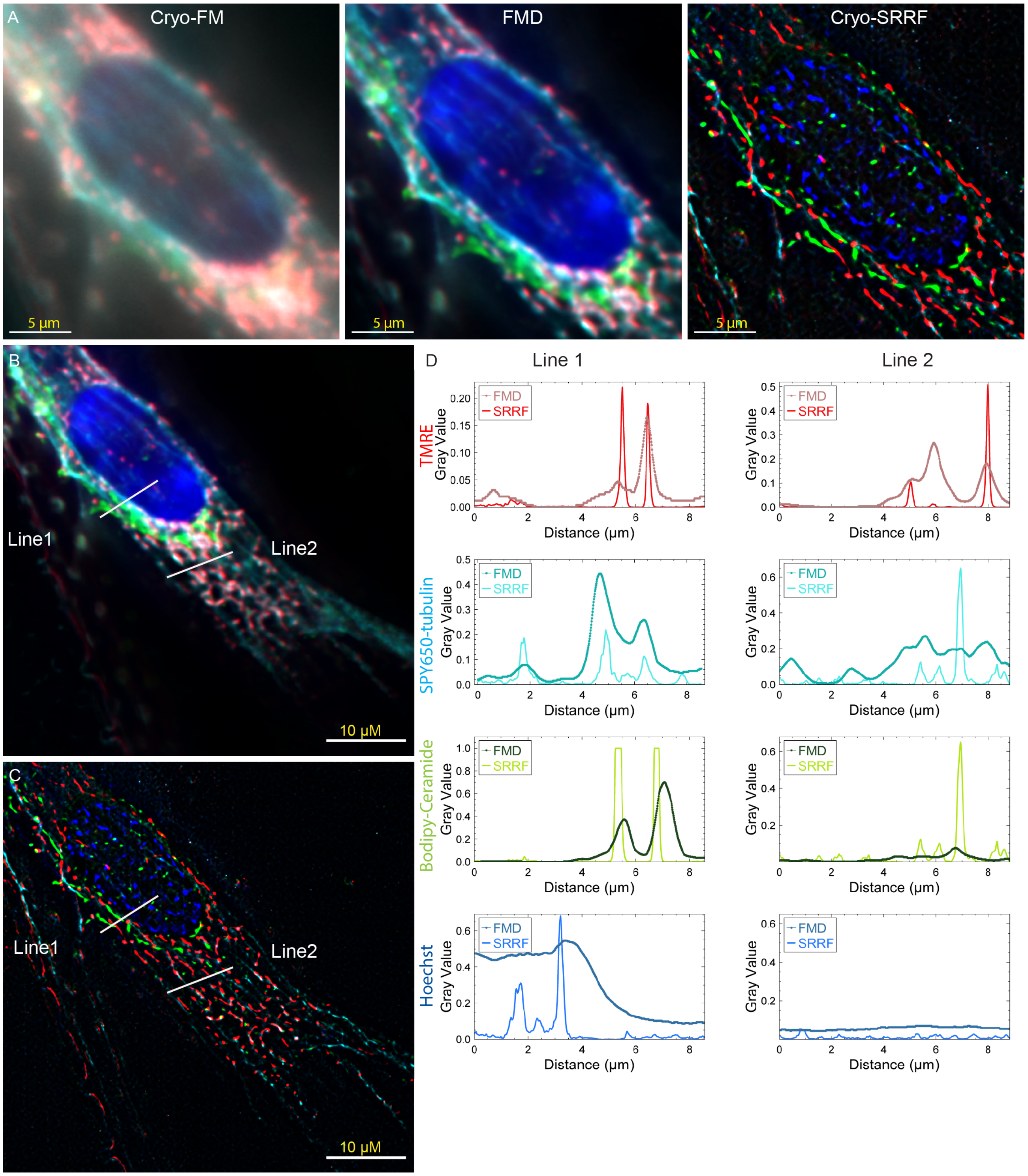


Fig S2: Comparing FMD to SRRF. (A) Comparing stabilized maxZ to FMD and cryo-SRRF (Fig 1). Profiles of two lines in the (B) FMD and (C) SRRF image are compared. The scale bar is 10 µM. (D) The plots of the profile of the FMD (filled circles connected by line) and SRRF (line). The TMRE, SIR-tubulin, Bodipy-Ceramide and Hoechst signal are represented by shades of red, cyan, green and blue, respectively.

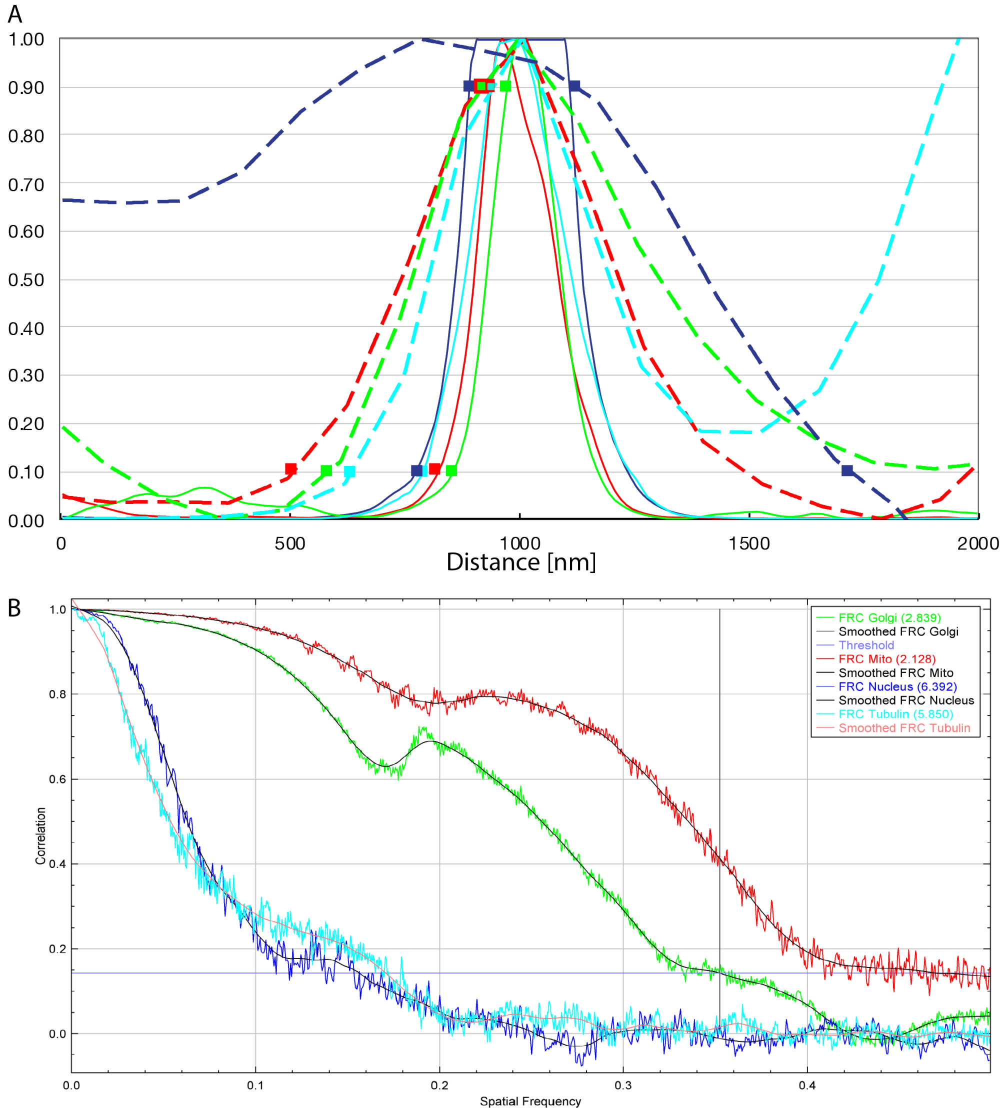


Fig S3: Resolution estimates: (A) Edge response of the FMD and cryo-SRRF analysis, and (B) FRC plots of the cryo-SRRF analysis. (A) Blue, red, green, and cyan lines represent one example (out of four) of line profiles of the Hoechst, TMRE, bodipy-ceramide, and SPY650-tubulin (respectively) staining from the FMD (dashed thicker lines) and cryo-SRRF (solid thinner lines). The squares represent 10 and 90 % values of the peaks in the line profiles. (B) Blue, red, green, and cyan lines represents the FRC values from the Hoechst, TMRE, bodipy-ceramide, and SPY650-tubulin staining, and the underlying black line represents the smoothed FRC signals, respectively. The threshold of 1/7 is represented in purple. The spatial frequencies at 1/7 threshold listed in the legend box and in table S1 are represented in units 1/Nyquist, and in Table S1 in nm.

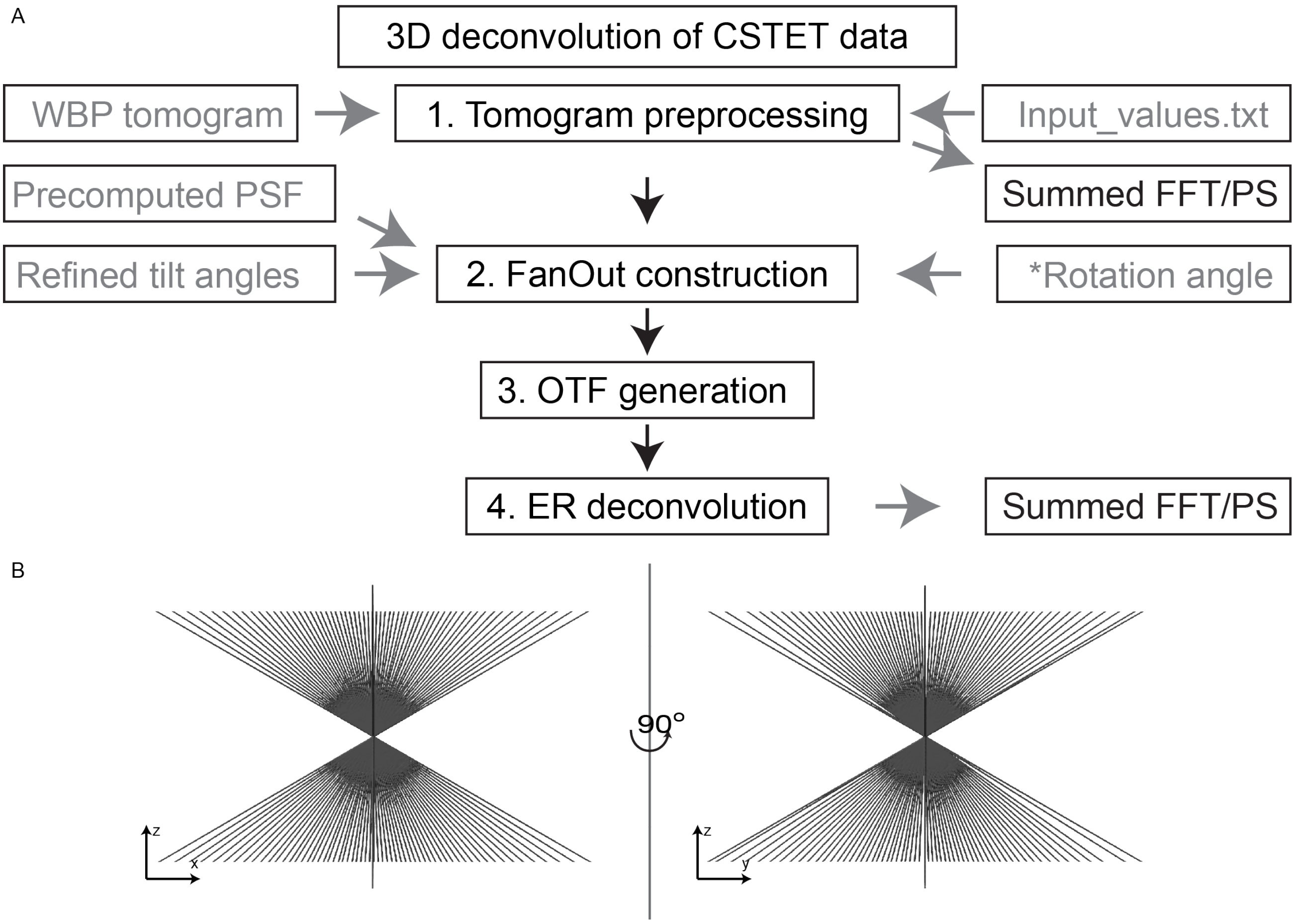


Fig S4: Workflow of the decon-script, and representation of a dual-axis multi-probe PSF: (A) The four steps of deconvolution and their respective inputs (grey) are depicted. Optionally, a summed FFT or PS can be computed from the input tomogram and the deconvolved data. (*) The Rotation angle is only needed for dual-axis tomogram. (B) A representative dual-axis multi-probe PSF is shown.

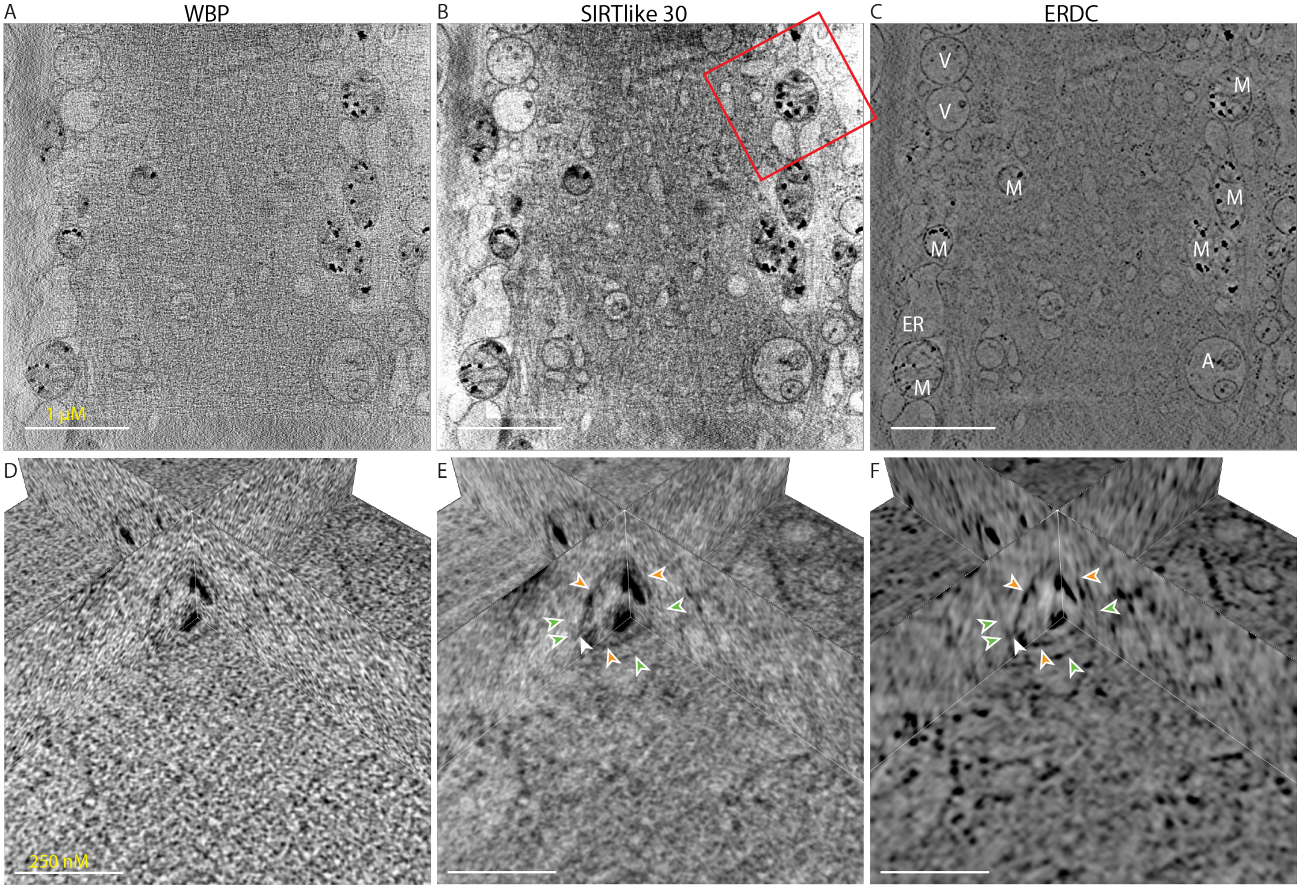


Fig S5: A deconvolved dual-axis tomogram in a different cell on the same grid around 750 nm thick. (A-C) XY slices for overview, (D-F) Orthoslice view of the box indicated in (B), (A,D) WBP tomogram (input for ERDC), (B,E) SIRTlike30-filtered, and (C,F) deconvolved tomogram. Many cellular components, such as vesicles (V), mitochondria (M), ER and autophagosomes (A) can be observed. In the orthoplanes, ERDC reveals additional details not easily seen in the WBP/SIRTlike30, such as an ER membrane (green arrowhead) surrounding a mitochondrion (OMM, orange arrowhead), or a protein density connecting the two membranes (white arrowhead). Scale bars in the overview and orthoplanes are 1 μm and 250 nm, respectively.

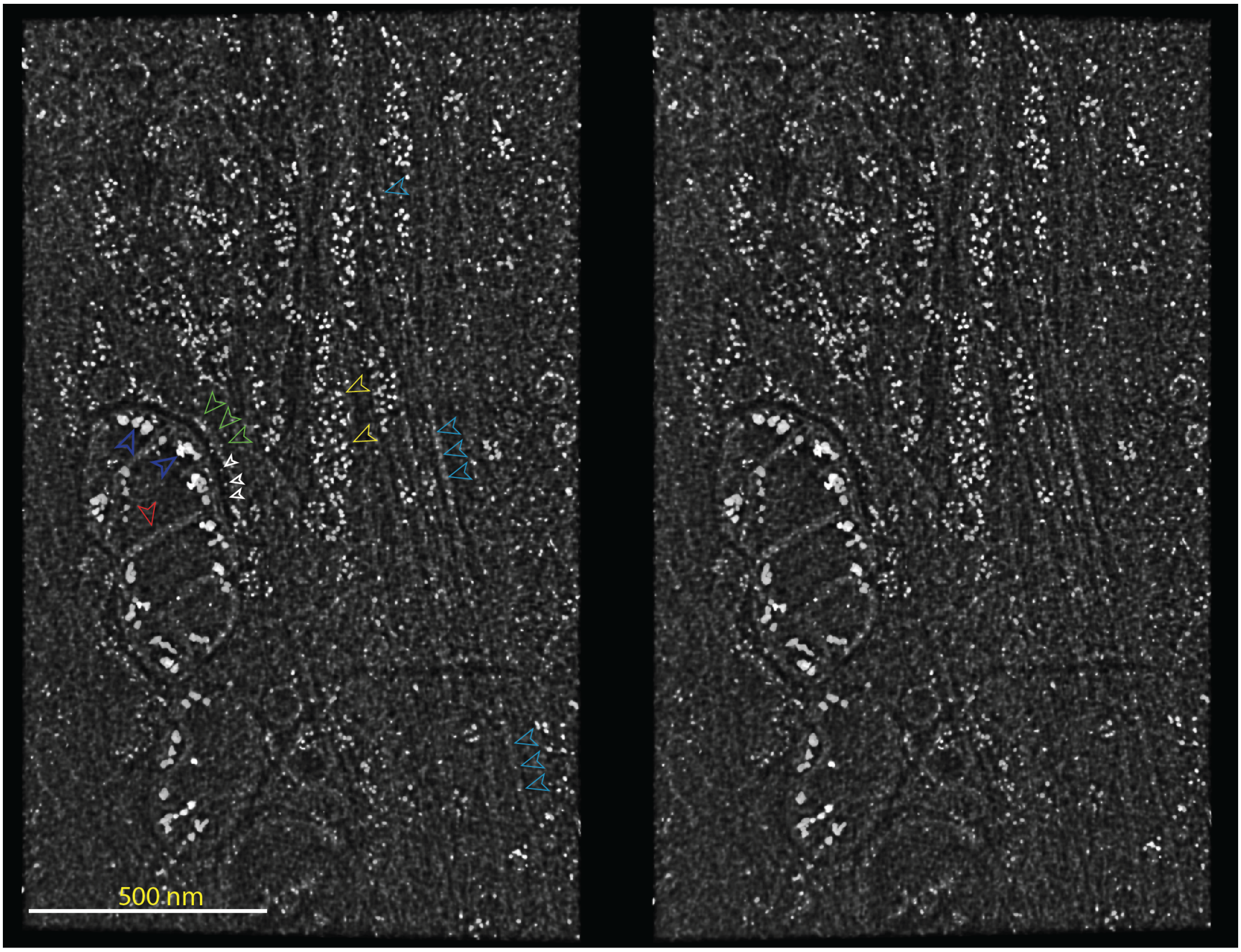


Fig S6: Stereo images of a 120 nm thick section of the same region as in Fig 2. Cristae (red arrowheads), CaP deposits (blue arrowheads), ER membrane (green arrowheads), ribosomes (yellow arrowheads), microtubules (cyan arrowheads), and proteins in the mitochondria-ER contact site are observable (white arrowheads). Slices 40-70 are depicted. Total region thickness is 850 nm.

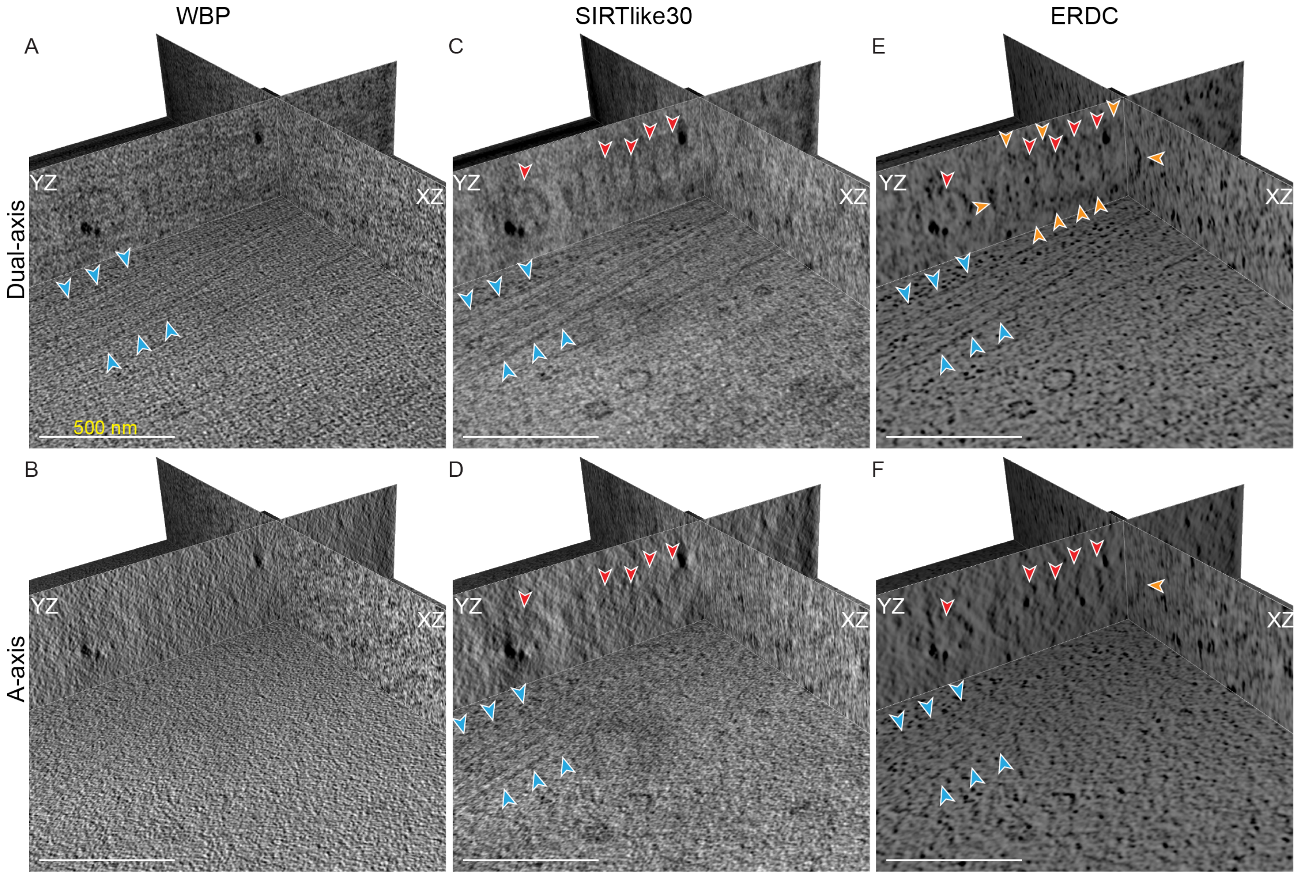


Fig S7: Orthoplanes of same area as in Fig 4*B,D,F*): Comparing dual- (A,C,E) to single-axis (B,D,F). (A,B) WBP tomograms. With the exception of some CaP (dark spots in the YZ image), the WBP shows little information. Some microtubules (cyan arrowheads) can be identified in the dual-axis tomogram (A), but are barely discernible in the single-axis tomogram (B). (C,D) SIRTlike30 filtered tomograms. When applying a SIRTlike filter, cristea (red arrowheads) of the mitochondria are revealed in the dual-axis tomogram (C), but not in the single-axis tomogram (D). (E,F) ERDC tomograms. The dual-axis ERDC tomogram (E) additionally reveals the OMM of the mitochondria in all directions (orange arrowheads), as well as the cristae (red arrowhead). However, the Z-axis elongation artifact obscures their identification in the single axis tomogram (F). Scale bar is 500 nm in all images.

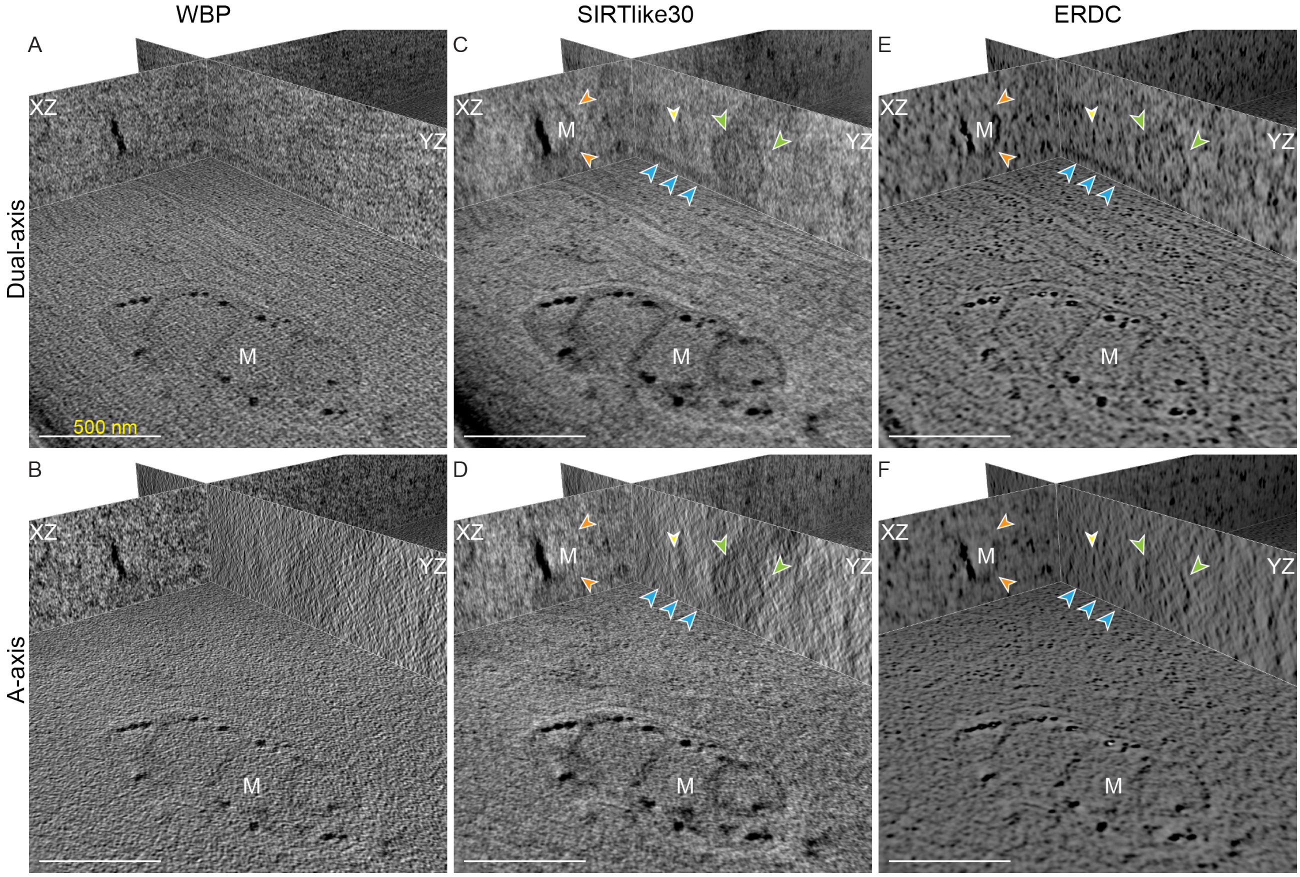


Fig S8: Orthoplanes of same area as in Fig 4*A,C,E*: Comparing dual- (A,C,E) to single-axis (B,D,F). (A,B) WBP tomograms. A mitochondrion (M) is observed in the XY plane, of the (A) dual-axis and (B) single-axis tomogram. (C,D) SIRTlike30 filtered tomograms. Dual-axis (C) reveals several more details such as a vesicle (green arrowheads), a microtubule running across the YZ plane (cyan arrowheads), and the OMM of a mitochondria (orange arrowheads), all of which are hard to identify in the single-axis tomogram (D). (E,F) ERDC tomograms. The dual-axis ERDC tomogram (E), reveals many more details, specially in the XZ, YZ planes. Many of these details are missing in the single-axis tomogram (F). Scale bars are 500 nm in all images.

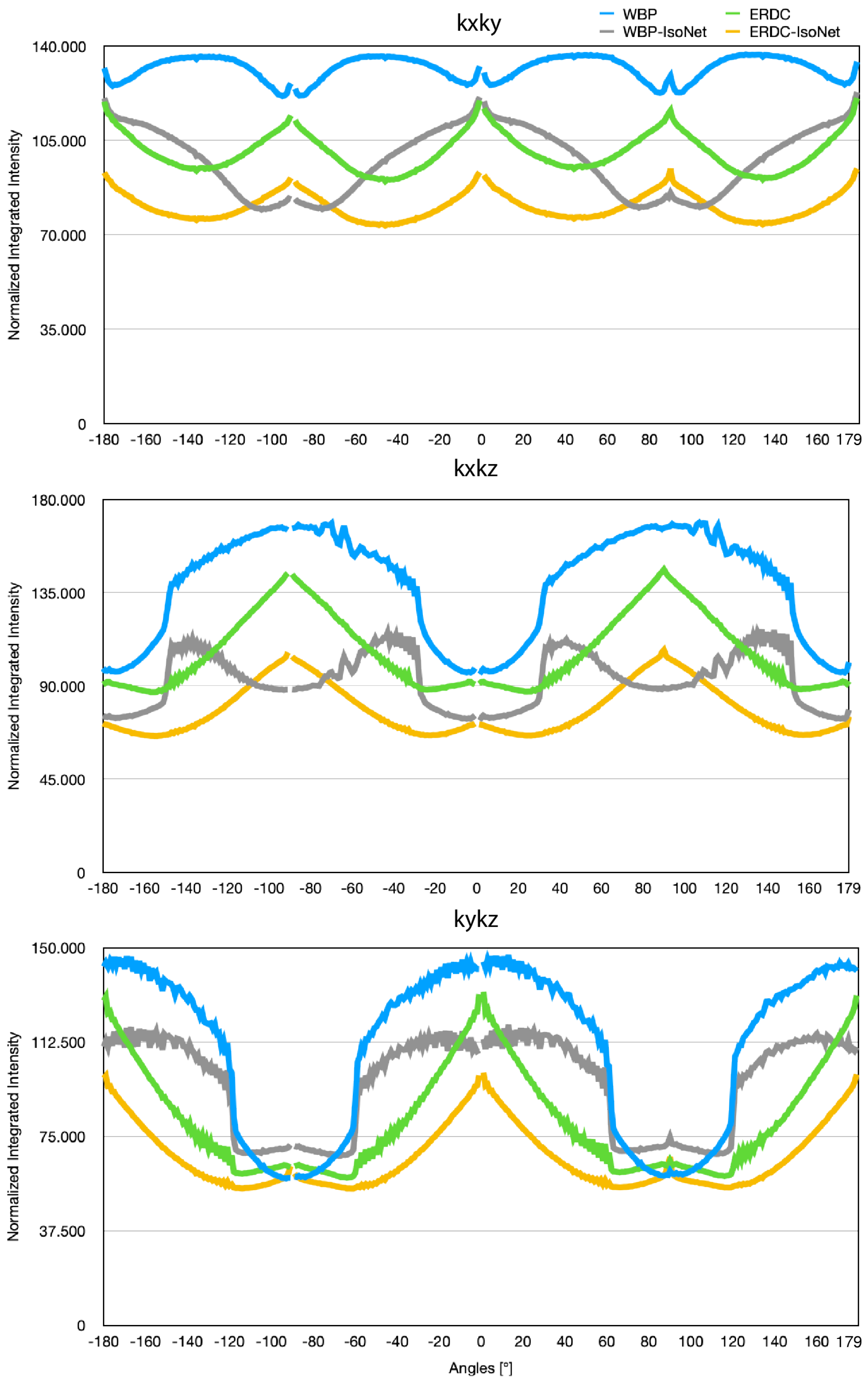


Fig S9: Azimuthal average of the power spectra from Fig 6.

### References

[1] David Kirchenbuechler, Yael Mutsafi, Ben Horowitz, Smadar Levin-Zaidman, Deborah Fass, Sharon G. Wolf, and Michael Elbaum. Cryo-STEM Tomography of Intact Vitrified Fibroblasts. *AIMS Biophysics*, 2(3):259–273, 2015. ISSN 2377-9098. doi: 10.3934/biophy.2015.3.259. URL `http://www.aimspress.com/article/10.3934/biophy.2015.3.259`.

[2] Johannes Schindelin, Ignacio Arganda-Carreras, Erwin Frise, Verena Kaynig, Mark Longair, Tobias Pietzsch, Stephan Preibisch, Curtis Rueden, Stephan Saalfeld, Benjamin Schmid, Jean-Yves Tinevez, Daniel James White, Volker Hartenstein, Kevin Eliceiri, Pavel Tomancak, and Albert Cardona. Fiji: an open-source platform for biological-image analysis. *Nature Methods*, 9(7):676–682, 7 2012. ISSN 1548-7091. doi: 10.1038/nmeth.2019. URL `http://www.nature.com/articles/nmeth.2019`.

[3] Nils Gustafsson, Siân Culley, George Ashdown, Dylan M. Owen, Pedro Matos Pereira, and Ricardo Henriques. Fast live-cell conventional fluorophore nanoscopy with ImageJ through super-resolution radial fluctuations. *Nature Communications*, 7:1–9, 2016. ISSN 20411723. doi: 10.1038/ncomms12471.

[4] Siân Culley, Kalina L. Tosheva, Pedro Matos Pereira, and Ricardo Henriques. SRRF: Universal live-cell super-resolution microscopy. *International Journal of Biochemistry and Cell Biology*, 101(March):74–79, 2018. ISSN 18785875. doi: 10.1016/j.biocel.2018.05.014. URL `https://doi.org/10.1016/j.biocel.2018.05.014`.

[5] Alex Herbert and Oliver Burri. Fourier Ring Correlation ImageJ Plugin, 2016. URL `https://github.com/BIOP/ijp-frc`.

[6] Robert P J Nieuwenhuizen, Keith A Lidke, Mark Bates, Daniela Leyton Puig, David Grünwald, Sjoerd Stallinga, and Bernd Rieger. Measuring image resolution in optical nanoscopy. *Nature Methods*, 10(6):557–562, 6 2013. ISSN 1548-7091. doi: 10.1038/nmeth.2448. URL `http://www.nature.com/articles/nmeth.2448`.

[7] Bob Dougherty. Diffraction PSF 3D, 2005. URL `https://www.optinav.info/Diffraction-PSF-3D.htm`.

[8] Cédric MessaoudiI, Thomas Boudier, Carlos Oscar Sanchez Sorzano, and Sergio Marco. TomoJ: tomography software for three-dimensional reconstruction in transmission electron microscopy. *BMC Bioinformatics*, 8(1):288, 12 2007. ISSN 1471-2105. doi: 10.1186/1471-2105-8-288. URL `https://bmcbioinformatics.biomedcentral.com/articles/10.1186/1471-2105-8-288`.

[9] Barnali Waugh, Sharon G. Wolf, Deborah Fass, Eric Branlund, Zvi Kam, John W. Sedat, and Michael Elbaum. Three-dimensional deconvolution processing for STEM cryotomography. *Proceedings of the National Academy of Sciences*, 117(44):27374–27380, 11 2020. ISSN 0027-8424. doi: 10.1073/PNAS.2000700117. URL `https://www.pnas.org/content/117/44/27374`.

[10] Hans Chen, Warren K. Clyborne, John W. Sedat, and David A. Agard. PRIISM: an integrated system for display and analysis of 3-D microscope images. In Raj S. Acharya,

Carol J. Cogswell, and Dmitry B. Goldgof, editors, *Biomedical Image Processing and Three-Dimensional Microscopy*, volume 1660, pages 784–790, 6 1992. doi: 10.1117/12.59604. URL http://proceedings.spiedigitallibrary.org/proceeding.aspx?articleid=987191.

[11] M. Arigovindan, J. C. Fung, D. Elnatan, V. Mennella, Y.-H. M. Chan, M. Pollard, E. Branlund, J. W. Sedat, and D. A. Agard. High-resolution restoration of 3D structures from widefield images with extreme low signal-to-noise-ratio. *Proceedings of the National Academy of Sciences*, 110(43):17344–17349, 10 2013. ISSN 0027-8424. doi: 10.1073/pnas.1315675110. URL http://www.pnas.org/cgi/doi/10.1073/pnas.1315675110.

[12] Philippe Carl. Azimuthal Average, 2007. URL https://imagej.nih.gov/ij/plugins/azimuthal-average.html.
